# Female reproductive fluid evolves rapidly to favor conspecific sperm

**DOI:** 10.64898/2026.05.14.725137

**Authors:** Livia Pinzoni, Elisa Morbiato, Owen C. Dorsey, Johaira Hernandez Melo, Alessandro Devigili, Clelia Gasparini, Gil G. Rosenthal

## Abstract

Avoiding fertilization with genetically incompatible partners, whether too similar or too divergent, is a central challenge for sexually reproducing organisms. Selection can favor mechanisms acting before and after mating, with postmating processes potentially compensating for constraints on premating choice. In the postmating context, female reproductive fluid (FRF) can modulate sperm performance and bias fertilization outcomes, but its contribution to reproductive isolation remains unclear. We tested whether FRF mediates discrimination against heterospecific and related sperm in two naturally hybridizing sister species of swordtails, *Xiphophorus birchmanni* and *X. malinche,* that diverge in premating behavior towards heterospecifics. Effects of FRF differed sharply between species. In *X. malinche*, FRF enhanced the velocity of conspecific sperm relative to heterospecifics, consistent with postmating discrimination against hybridization. In contrast, FRF in *X. birchmanni* did not favor conspecific sperm. Evidence for inbreeding avoidance was weaker, and we found no indication of a trade-off between discrimination against genetically similar and dissimilar sperm. These results show that female reproductive fluid can serve as a rapidly evolving axis of reproductive isolation through postmating female choice.

## Introduction

Successful reproduction and offspring fitness depend on the compatibility of interacting gametes. From an evolutionary perspective, individuals are therefore expected to avoid fertilizations involving gametes that are either too genetically similar, increasing the risk of inbreeding depression, or too genetically divergent, resulting in outbreeding depression (1). Selection for genetic compatibility can act at multiple stages of reproduction. While mate choice before mating can bias mating success, the opportunity for choice does not end with copulation. Females can also exert cryptic choice (2,3) after mating, through physiological and biochemical processes that influence gamete interactions at or beyond fertilization. As before mating, postmating female choice often favors conspecific sperm over heterospecific sperm, a pattern known as conspecific sperm precedence (4). At the other end of the spectrum, post-pollination processes reliably prevent fertilization by genetically similar gametes in plants (5,6), while few studies have evidence for postmating avoidance of kin in animals (but see 7).

Cryptic female choice is expected to be stronger when premating mate-choice mechanisms are limited or costly (8,2). Across species, conspecific preferences before mating are often context dependent, varying with the physical or social environment, and proximate constraints may limit opportunities for kin avoidance before mating (9). If females have less opportunity to choose before mating, postmating mechanisms are expected to enable them to bias fertilization outcomes and increase the likelihood of fertilization by compatible mates.

Female-derived reproductive fluid is increasingly recognized as an important mechanism of cryptic female choice via its differential effect on sperm performance across a wide range of externally and internally fertilizing species. The female reproductive fluid (FRF) is defined broadly as “the medium, arising from females, through which sperm must pass on their way to fertilize the eggs” (10) and accumulating evidence shows that FRF can differentially affect multiple components of sperm performance, including chemotaxis, velocity, trajectory, motility, and viability (reviewed in 10). In several taxa, FRF biases sperm performance in favor of conspecific sperm (11-15). In salmonids, for instance, FRF mediates cryptic female choice by altering sperm swimming behavior, promoting chemoattraction and fertilization success of conspecific sperm. Across systems, the strength of this species-specific discrimination appears to correlate with the risk of hybridization. FRF enhances conspecific sperm performance in *Ficedula* flycatchers, which commonly hybridize, but not in bird species with low hybridization risk (15,16). A comparable pattern occurs within species, where FRF favors sperm from a female’s own population over more distant populations (17). In guppies, a viviparous poeciliid fish, FRF not only favors males from a female’s own population (23) but also favors unrelated males over siblings (7). This suggests that FRF-mediated choice, like gamete recognition in flowering plants (19) may be able to simultaneously filter genetically similar and dissimilar mates, and that FRF effects should be stronger when premating choice is constrained (9).

We used a powerful experimental system to test for both conspecific sperm precedence and avoidance of inbreeding within the same individual females. The sister swordtail species *Xiphophorus birchmanni* and *X. malinche* (Teleostei: Poeciliidae) hybridize in nature; early-generation hybrids have reduced fitness due to genetic incompatibilities (20, 21), including melanoma (22) and, most acutely, mitonuclear dysfunction (23). Both species should thus be expected to avoid heterospecific fertilizations. Both species mate multiply in the wild, with most females carrying offspring from 2 or 3 sires (24, 25). This provides ample opportunity for postmating, pre-fertilization mate choice in the reproductive tract.

In swordtails, female premating choice relies on olfactory mechanisms (26, 27); interference with olfaction likely triggered ongoing (28) and continuing (29) hybridization between the two species. Mating preferences and individual behavior are divergent between the highland *X. malinche* and the lowland *X. birchmanni. X. birchmanni* females prefer conspecifics, reinforced by preferences for visual cues (30). Preferences are developed through social experience with conspecifics (31-33).

By contrast, in *X. malinche,* olfactory preferences are weaker and their expression is dependent on heterospecific (*X. birchmanni*) experience for activation (31, 27). Females *X. malinche* prefer visual cues of *X. birchmanni* over those of conspecific males. Overall, therefore, *X. malinche* have weaker premating preferences for conspecifics. Compared to *X. birchmanni,* individual *X. malinche* also show reduced exploratory behavior and increased time in shelter (33, 34), likely reducing the opportunity to engage in courtship interactions before mating. Overall premating choice is thus expected to be weaker in *X. malinche,* which should favor stronger postmating preferences.

We tested postmating preference by assessing how female reproductive fluid from individual females differentially influence traits linked to fertilization success of sperm from a sibling, an unrelated conspecific and a heterospecific male. Female *X. malinche,* but not *X. birchmanni,* accelerated the sperm of unrelated, conspecific males, showing strong conspecific sperm precedence and a moderate bias against sperm of full brothers.

## Materials and Methods

### Overview of Experimental design

We strove for a fully factorial, crossed repeated measures design to test the postmating preference of individual female *X. birchmanni* and *X. malinche* for sibling, unrelated conspecific, and heterospecific males through differential interactions between sperm and female reproductive fluid. The experimental design aimed to test each male sperm with FRF from a female representing each category (i.e., one sibling, one unrelated conspecific, and one heterospecific female). However, not all intended combinations were achieved due to practical constraints, short post-extraction viability of sperm bundles, and occasional failure of males to produce sperm (see supplementary information). Trials were balanced across species: for *X. birchmanni*, we performed 20 sibling trials (from 6 families), 20 unrelated trials, and 30 heterospecific trials using N=29 females and 39 males. For *X. malinche*, we performed 18 sibling trials (from 5 families), 18 unrelated trials, and 30 heterospecific trials, using N=30 females and 31 males.

We measured sperm curvilinear velocity (VCL), a standard proxy of fertilization success in fish (35, 36) and in poecilids (37, 38). We measured sperm velocity both in a control solution (0.9% NaCl) and in the FRF of sibling, unrelated conspecific, and heterospecific females (Fig. 1). This allowed us to compare the strength of FRF effects between species and across assays (conspecific preference and kin avoidance) and to test for evidence of a tradeoff between kin avoidance and conspecific preference. Upon collection of FRF samples, all females and males were photographed using a standard protocol (39) to allow for subsequent morphological analysis. See Supplementary Information for details on compliance, husbandry, generation of sibling cohorts and ethic permit.

**Figure 1.**
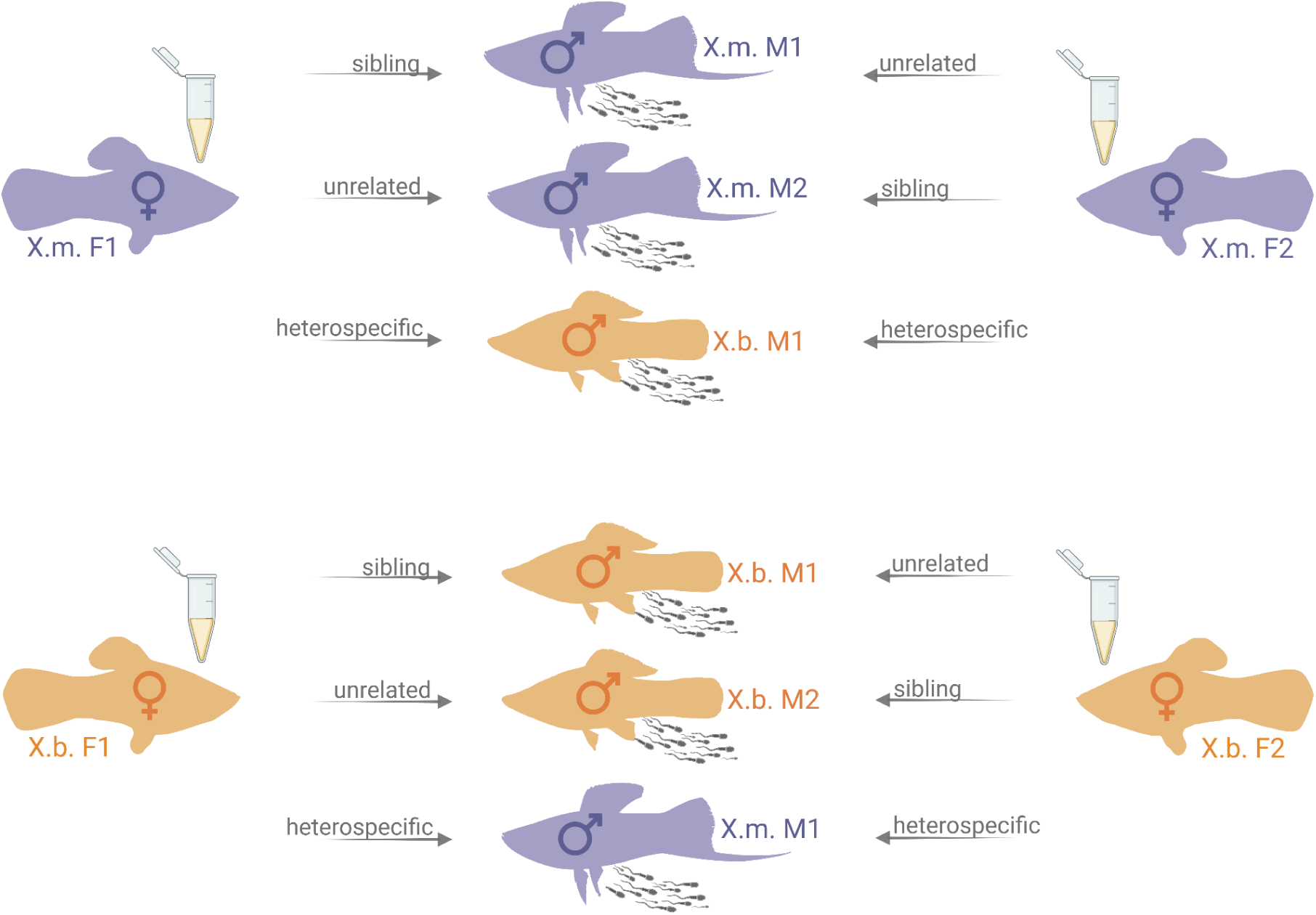
Experimental design for assessing effects of female reproductive fluid (FRF) on sperm performance in the *Xiphophorus malinche* – *X. birchmanni* system. FRF was collected from females of each species (*X. malinche* in purple, *X. birchmanni* in orange) and used to test its effect on sperm performance in three male categories: sibling, unrelated conspecific, or heterospecific. M1, M2 = male 1, male 2; F1, F2 = female 1, female 2.

### Collection of FRF and ejaculate

At least three days prior to the experiment, individuals were housed in same-sex groups to ensure that FRF was free of sperm and that males produced a sufficient quantity of spermatozoa. Female reproductive fluid was collected using methods from Gasparini & Pilastro (7). Briefly, females were anesthetized in a water bath containing buffered tricaine methanesulfate (MS-222, 125–150 mg L⁻¹) and positioned belly-up on a wet sponge under a stereomicroscope. After anesthesia, 3 µl of 0.9% NaCl was carefully injected into the female gonoduct using a Drummond micropipette and then retrieved. This process was repeated 4–5 times, resulting in an FRF sample of approximately 12–15 µl per female. Each FRF sample was divided into three aliquots, used to evaluate sperm velocity for one sibling, one unrelated conspecific, and one heterospecific male in a randomized order.

Sperm were collected from anesthetized males positioned as described for females, adapting the protocol used in zebrafish (40) to our study system. Briefly, males were anesthetized in the water bath containing MS-222 and positioned belly-up on a wet sponge under a stereomicroscope. Gentle abdominal pressure was applied in a head-to-tail direction near the gonopore, which was positioned against a glass capillary tube. The ejaculate was drawn into the tube via capillary action and immediately transferred to a 20 µL drop of saline solution (0.9% NaCl) placed on a black microscope slide to enhance visual contrast of the sperm bundles.

### Assessment of sperm performance

Methods followed (7). For each female, we assayed sperm in a solution of 60% (v/v) FRF with 40% of KCl (150 mM) and 4 mg ml⁻¹ of BSA. We sought to test each female with sperm of (a) a sibling (b) an unrelated conspecific male, and (c) a heterospecific male. Six intact sperm bundles (in swordtails sperm are packaged into discrete units known as spermatozeugmata or bundles), from each male (n = 70, 39 *X. birchmanni* and 31 *X. malinche* males) were placed under a compound microscope (Ernst Leitz Wetzlar, Germany) in separate wells on a multi-well slide containing one of the three medium with FRF (a, b, c) as well as a control solution containing 0.9% NaCl instead of FRF.

Thirty-second videos of sperm motility were recorded (in random order) using an iPhone HD camera with High Dynamic Range (HDR) settings at 60 frames per second. Videos were subsequently analyzed to extract curvilinear velocity (VCL, µm s⁻¹) of sperm spontaneously leaving the bundle. At least three out of the six bundles per male were analyzed (mean number of motile sperm per analysis: 162.9 ± 44.1 s.d.). All analyses were conducted using a CEROS sperm tracker (Hamilton-Thorne Research, Beverly, MA, USA).

To assess the repeatability of sperm performance measures, we conducted additional assays using a separate sample of 15 males and 15 females per species drawn from stock populations of unknown relatedness. For each of those males, sperm velocity was measured twice per condition using the same ejaculate: two independent activations and video recordings were performed in control solution and two in the FRF of the same conspecific female (technical replicates within male–female combinations).

Sperm performance was highly repeatable in both environments (control solution: r = 0.983; conspecific FRF: r = 0.950; all p<0.001; Supplementary Material), indicating consistent FRF effects on sperm velocity within male–female combinations.

### Statistical analysis

To estimate the effects of FRF as a mate-choice mechanism, we firstly calculated the change in percentage of curvilinear velocity (VCL), between sperm cells swimming in the control solution and in each FRF type: sibling, unrelated conspecific, and heterospecific for all males (sample size N = 39 *X. birchmanni*; N = 31 *X. malinche*). We built linear mixed effect (LME) models using the Maximum Likelihood (ML) fit method with a Gaussian error distribution to estimate the probability of sperm performance variation associated with the FRF treatment (sibling, unrelated conspecific, heterospecific), the species (*X. birchmanni* – *X. malinche*), and their interaction. In model 1 we set VCL change as the dependent variable and we used the FRF type, the species, and their interaction as fixed effects. To investigate the relationship between male and female morphology and VCL, we also included male and female body length as predictors. Finally, we incorporated the residual of body depth onto standard length (hereafter fineness index) as a proxy for female reproductive state: low residuals indicate poor-condition or nonreproductive females, while high residuals indicate gravid females in the later stages of pregnancy (41). We included females’ morphology for analysis because female poeciliids can store sperm for several months before fertilization (42) and swordtails exhibit proceptive behavior towards males throughout the reproductive cycle (43).

We included in our model male and female identities as random factors to account for effects of individual identity on sperm performance variation and the non-independence of the data collected from the same males and females. We ran a full model comprehensive of all factors and their interactions, and then removed non-significant terms starting with the highest *p-*value to retain in the final model only significant factors and interactions (Supplemental Information Model 2 SI; Model 3 SI). Linear mixed-effects models were compared using Akaike’s Information Criterion (AIC) to evaluate relative model support, and the model with the lowest AIC was selected for inference.

Because sperm velocities in the control solution differed significantly between *X. malinche* and *X. birchmanni* (t-test p = 0.001, supplementary table S2), and because species significantly interacted with fluid type in the global Model 1 (species × fluid type p < 0.05), we also present species-specific linear mixed-effects models. To elucidate the direction of VCL change, we included the fixed effects control VCL, fluid type, and their interaction, and random intercepts for male ID and female ID.

Fixed effects were evaluated using model summary. Post-hoc pairwise comparisons among the levels of fluid type were performed using linear contrasts on the fixed effects of the fitted linear mixed-effects models (LME) for each species. Contrasts were constructed to compare each pair of levels while accounting for covariates and random effects. P-values were calculated using MATLAB’s coefficient test function.

To assess models adequacy, residual diagnostics were performed, including plots of residuals versus fitted values, QQ-plots, residual histograms, and comparisons of residuals across treatment groups.

Furthermore, to assess species-specific differences in sperm velocity, we compared the change VCL between *X. birchmanni* and *X. malinche* within each fluid environment (sibling, unrelated conspecific, heterospecific). For each environment, Cohen’s d was calculated as the difference between the mean VCL of *X. malinche* and the mean VCL of *X. birchmanni*, divided by the pooled standard deviation of the two species. The pooled standard deviation was computed using the standard formula for two independent groups. Cohen’s *d* was calculated for each comparison to quantify the magnitude of species differences, following the standard formulation for two independent samples (44).

To test for potential trade-offs between avoidance of heterospecific and related sperm, we computed individual-level ΔVCL values per male–female pair for each treatment, and used Pearson correlations to assess covariation among the three preference dimensions (sibling, unrelated conspecific, heterospecific). Correlations were computed separately for each species.

Since sperm (45) and FRF traits may show heritable variation within species, we controlled for any effect of the family identity on VCL variation by running a LME model with VCL change as the dependent variable, male family as predictor and male and female identity as random intercepts to account for repeated measures across trials.

Analyses were run in R version 4.4.2 and MATLAB R2024a.

## Results

### Overall model

We evaluated the magnitude of FRF effects on sperm performance as a function of male and female species identity and relatedness (model 1, Table 1). The model shows a significant interaction between species and the fluid type. Reproductive fluid of female *X. malinche* boosted sperm velocity of both sibling and unrelated conspecific males relative to heterospecifics. In contrast, female *X. birchmanni* FRF showed no differential effect across sperm types (Figure 2). Since neither male nor female standard length nor fineness index predicted VCL change, we removed these predictors from the analysis; results of intermediate models are presented in the Supplementary Information (Model 2, SI Table 3; Model 3, SI Table 4, Model 4, SI Table 5). Model comparison based on Akaike’s Information Criterion (AIC) indicated that the full model (Model 1) including all covariates and interactions provided the best fit to the data, with substantially lower AIC (1177.4) than any of the reduced models (AIC 1268–1269).

**Table 1.**
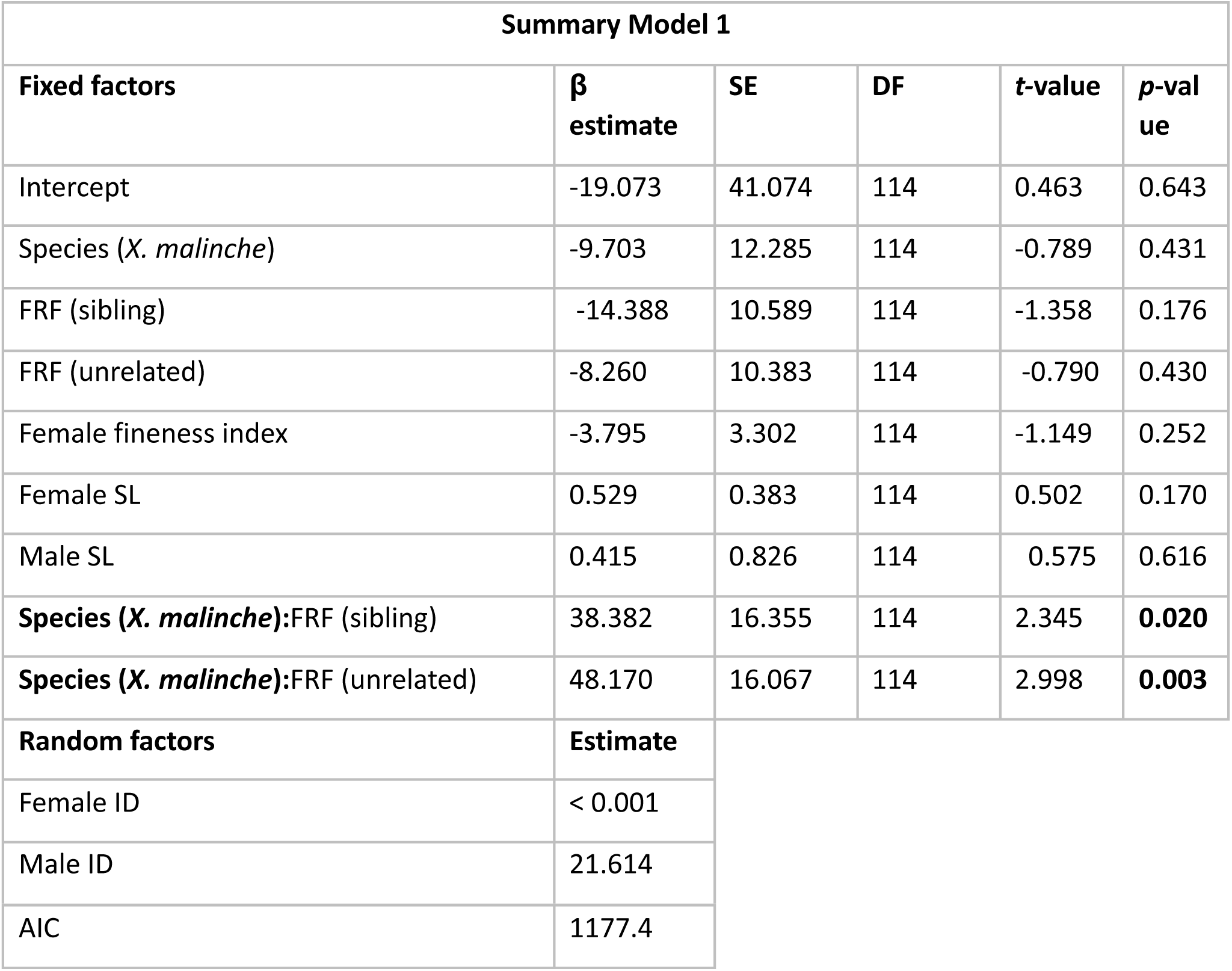
Results of LME (Model 1) including VCL change as dependent variable; male SL, female SL, female fineness index (residuals), species, FRF type, and their interaction as fixed factors; and female and male identity as random factors. Significant terms are in bold. Regression coefficients of the interactions refer to the *X. malinche* species while *X. birchmanni* was set to 0. FRF type was treated as a categorical variable with heterospecific as the reference level. SE=Standard Error; DF=Degrees of Freedom.

| Summary Model 1 |  |  |  |  |  |
| --- | --- | --- | --- | --- | --- |
| Fixed factors | $\beta$<br>estimate | SE | DF | t-value | p-value |
| Intercept | -19.073 | 41.074 | 114 | 0.463 | 0.643 |
| Species (*X. malinche*) | -9.703 | 12.285 | 114 | -0.789 | 0.431 |
| FRF (sibling) | -14.388 | 10.589 | 114 | -1.358 | 0.176 |
| FRF (unrelated) | -8.260 | 10.383 | 114 | -0.790 | 0.430 |
| Female fineness index | -3.795 | 3.302 | 114 | -1.149 | 0.252 |
| Female SL | 0.529 | 0.383 | 114 | 0.502 | 0.170 |
| Male SL | 0.415 | 0.826 | 114 | 0.575 | 0.616 |
| **Species (*X. malinche*):FRF (sibling)** | 38.382 | 16.355 | 114 | 2.345 | **0.020** |
| **Species (*X. malinche*):FRF (unrelated)** | 48.170 | 16.067 | 114 | 2.998 | **0.003** |
| Random factors | Estimate |  |  |  |  |
| Female ID | < 0.001 |  |  |  |  |
| Male ID | 21.614 |  |  |  |  |

|  |  |
| --- | --- |
| AIC | 1177.4 |

**Figure 2.**
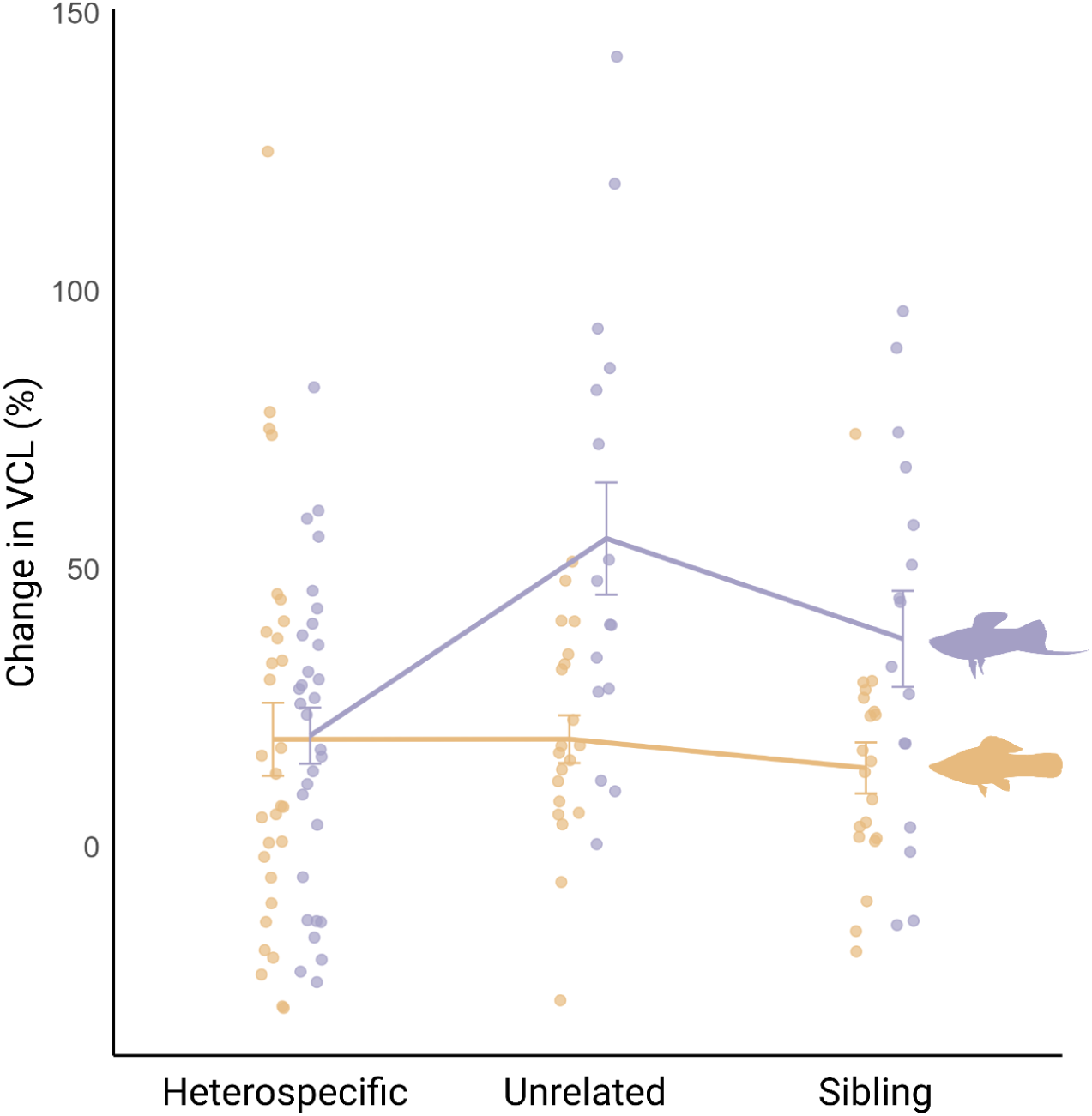
Change in sperm curvilinear velocity (VCL) from control solution to female reproductive fluid (FRF) across FRF types for *X. birchmanni* (orange) and *X. malinche* (purple). Points represent raw data of individual measurements of VCL change. Lines connect species-specific mean values across FRF types, with error bars indicating ± standard error of the mean. The plot highlights species-specific patterns in FRF effects on sperm performance, with *X. malinche* (purple) showing stronger differentiation among FRF types compared to *X. birchmanni* (orange), consistent with results from mixed-effects models (Tables 1, 2 and 3).

Given that species significantly interacted with FRF type in the global Model 1 ( species × FRF type p < 0.020; p < 0.003), and since control sperm velocities differed significantly between *X. malinche* and *X. birchmanni* (t-test *p* < 0.001, Supplemental Information), we ran species-specific linear mixed-effects models for FRF effects on sperm performance. In order to elucidate the direction of VCL change, we included the fixed effects control VCL, fluid type, and their interaction, and random intercepts for male ID and female ID.

### Species-specific models of FRF effects on sperm performance

We ran separate analyses for *X. malinche* sperm and *X. birchmanni* FRF. In *X. malinche* (Model 2; Table 2; Figure 3a), sperm velocity increased significantly in conspecific (unrelated) female fluid relative to heterospecific fluid (β = +139.4 ± 38.6, *p* < 0.001), while the effect of sibling fluid was positive but not significant (*p* = 0.11). The enhancement was strongest for males whose sperm was slower in the control solution (negative interaction between control VCL and conspecific fluid, β = −1.22 ± 0.39, *p* < 0.001; Table 2; Figure 3a).

**Table 2.** Results of LME for *X. malinche* sperm, including VCL change as dependent variable; VCL control, FRF type, and their interaction as fixed factors, and female and male identity as random factors. FRF type was treated as a categorical variable with heterospecific as the reference level. Significant terms are in bold. SE=Standard Error; DF=Degrees of Freedom.

| Summary Model 2 *X. malinche* |  |  |  |  |  |
| --- | --- | --- | --- | --- | --- |
| Fixed factors | $\beta$ estimate | SE | DF | t-value | p-value |
| Intercept | 106.63 | 18.844 | 56 | 5.658 | < 0.001 |
| FRF (Sibling) | 62.413 | 38.629 | 56 | 1.615 | 0.111 |
| **FRF (Unrelated)** | **139.39** | 38.629 | 56 | 3.608 | **< 0.001** |
| **VCL control** | **-0.785** | 0.166 | 56 | -4.706 | **< 0.001** |
| VCL control:FRF (Sibling) | -0.603 | 0.388 | 56 | -1.552 | 0.126 |
| **VCL control:FRF (Unrelated)** | **-1.224** | 0.388 | 56 | -3.152 | **0.002** |
| Random factors | Estimate |  |  |  |  |
| Female ID | < 0.001 |  |  |  |  |
| Male ID | < 0.001 |  |  |  |  |

**Figure 3.**
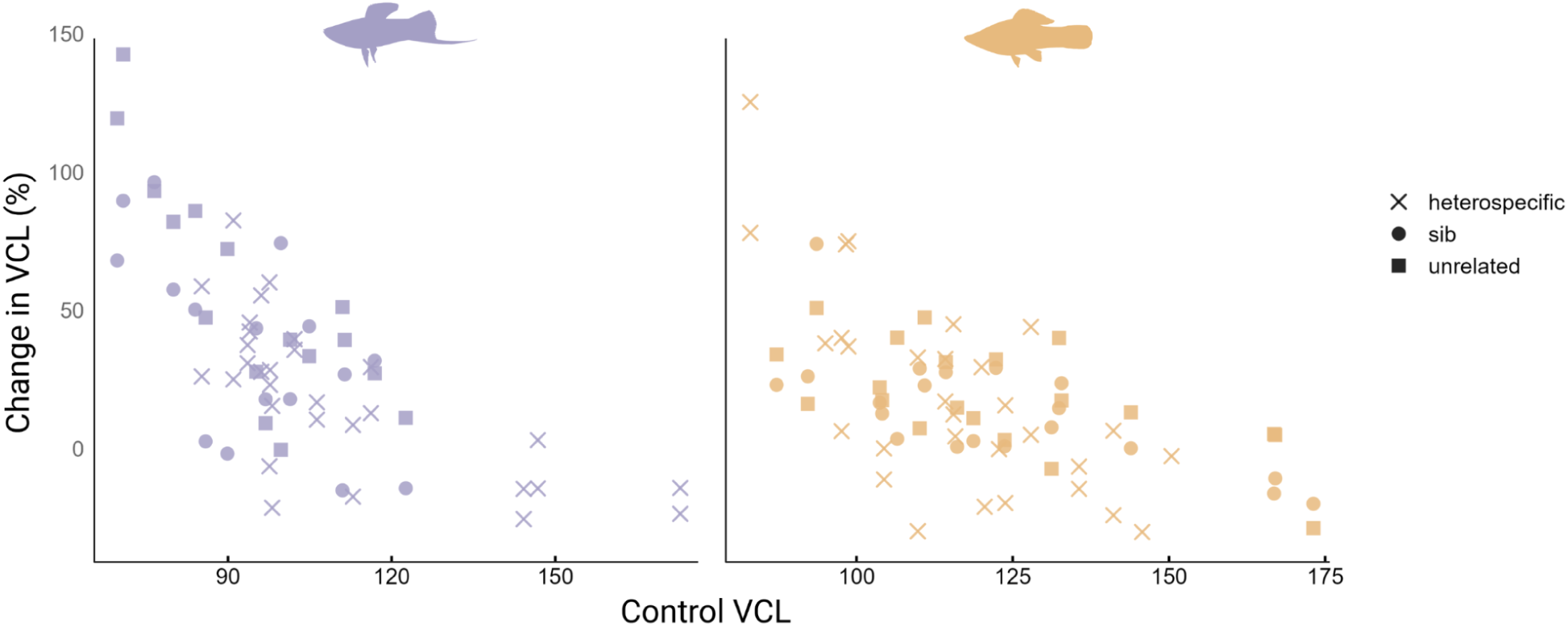
Scatter plot showing the relationship between control VCL and VCL change for *X. malinche* (purple) and *X. birchmanni* (orange). Symbols indicate fluid type: circles for sibling, squares for unrelated conspecific and asterisks for heterospecific. The plot highlights the relationship between VCL change and control VCL and the interaction with FRF type, consistent with the results from the mixed-effects models (Table 2, Table 4).

Accordingly, pairwise contrasts revealed that sperm performance differed significantly between heterospecific and unrelated conspecific fluids (p < 0.001, Table 3), while heterospecific vs sibling and sibling vs unrelated comparisons were not significant (p > 0.1, Table 3).

**Table 3.** Pairwise post-hoc comparisons among FRF levels in *X. malinche* from the species-specific linear mixed-effects model. P-values were obtained using linear contrasts on fixed effects, accounting for covariates and random effects included in the model.

| Pairwise Comparison | p-value |
| --- | --- |
| Heterospecific vs Sib | 0.1117 |
| **Heterospecific vs Unrelated** | **6.50e-4** |
| Sib vs Unrelated | 0.1114 |

In contrast, *X. birchmanni* (Model 3; Table 4; Figure 3b) sperm exhibited the opposite pattern. Sperm were slower in *X. birchmanni* fluid than in heterospecific fluid, for both unrelated conspecific (β = −116.2 ± 26.2, *p* < 0.001) and sibling fluids (β = −109.2 ± 26.2, *p* < 0.001). However, positive interactions between control VCL and both unrelated conspecific and sibling fluids (β = +1.04 ± 0.22, *p* < 0.001; β = +0.94 ± 0.22, *p* < 0.001, respectively) revealed that this effect was smaller in faster ejaculates. Consequently, *X. birchmanni* sperm performed worse in their own species’ fluids, especially if they were slower in the control solution.

**Table 4.** Results of LME (*X. birchmanni*) including VCL change as dependent variable; VCL control, FRF type, and their interaction as fixed factors, and female and male identity as random factors. FRF type was treated as a categorical variable with heterospecific as the reference level. Significant terms are in bold. SE=Standard Error; DF=Degrees of Freedom.

| Summary Model 3 *X.birchmanni* |  |  |  |  |  |
| --- | --- | --- | --- | --- | --- |
| Fixed factors | $\beta$ estimate | SE | DF | t-value | p-value |
| Intercept | 198.93 | 20.392 | 64 | 9.755 | < 0.001 |
| **FRF (Sibling)** | **-109.22** | 26.196 | 64 | -4.169 | **< 0.001** |
| **FRF (Unrelated)** | **-116.23** | 26.196 | 64 | -4.437 | **< 0.001** |
| **VCL control** | **-1.556** | 0.172 | 64 | -9.006 | **< 0.001** |
| **VCL control:FRF (Sibling)** | **0.939** | 0.215 | 64 | 4.354 | **< 0.001** |
| **VCL control:FRF (Unrelated)** | **1.035** | 0.215 | 64 | 4.799 | **< 0.001** |
| **Random factors** | **Estimate** |  |  |  |  |
| Female ID | 18.751 |  |  |  |  |
| Male ID | 4.0722 |  |  |  |  |

Pairwise contrasts further showed that sperm performance differed significantly between heterospecific and both sibling and unrelated conspecific fluids (p < 0.001 for both comparisons, Table 5), whereas no significant difference was detected between sibling and unrelated conspecific fluids (p = 0.74, Table 5).

**Table 5.**
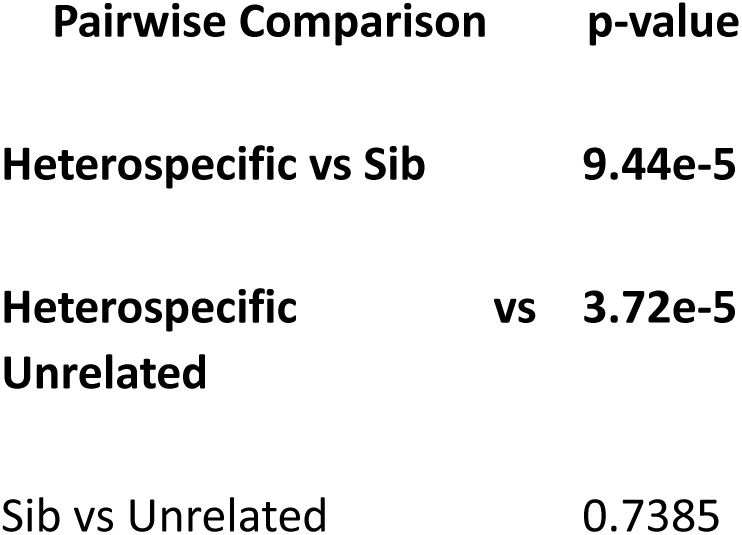
Pairwise post-hoc comparisons among FRF levels in *X. birchmanni* from the species-specific linear mixed-effects model. P-values were obtained using linear contrasts on fixed effects, accounting for covariates and random effects included in the model.

| Pairwise Comparison | p-value |
| --- | --- |
| **Heterospecific vs Sib** | **9.44e-5** |
| **Heterospecific vs Unrelated** | **3.72e-5** |
| Sib vs Unrelated | 0.7385 |

Overall, these results reveal a striking species asymmetry: *X. malinche* FRF favors sperm from conspecific, unrelated males, while *X. birchmanni* FRF slowed sperm velocity in both sibling and unrelated conspecific mates. The direction and magnitude of these effects depended on the initial sperm velocity, suggesting differences in how sperm–fluid compatibility and ejaculate performance interact between the two species.

### Effect sizes of female species on FRF responses across sperm types

The magnitude of species-specific differences in VCL change varied across FRF types: in heterospecific fluid, Cohen’s d = 0.016, 95% CI [–0.05, 0.08]) indicates a negligible difference in VCL change between species: heterospecific sperm were similarly slow in both *X. birchmanni* and *X. malinche* FRF. Female *X. malinche* boosted the velocity of both siblings (d = 0.841, 95% CI [0.70, 0.98]) and unrelated conspecifics (d = 1.187, 95% CI [0.95, 1.42]) substantially more than *X. birchmanni*.

Overall, species-specific differences in FRF effects on sperm motility were negligible in heterospecific fluid, large in sibling fluid, and very large in unrelated conspecific fluid, highlighting the role of fluid relatedness in modulating interspecific variation in sperm performance.

### Covariation in FRF effects across male types

To assess whether female reproductive fluid (FRF) exert consistent or independent effects on sperm from different male types, we examined pairwise Pearson correlations among ΔVCL values across the three treatments (sibling, unrelated conspecific, and heterospecific males). Because not all females were tested with all three male types, correlations were computed separately for each pairwise comparison and species. P-values were adjusted for multiple comparisons (three tests per species) using Holm’s correction (Fig. 4).

**Figure 4.**
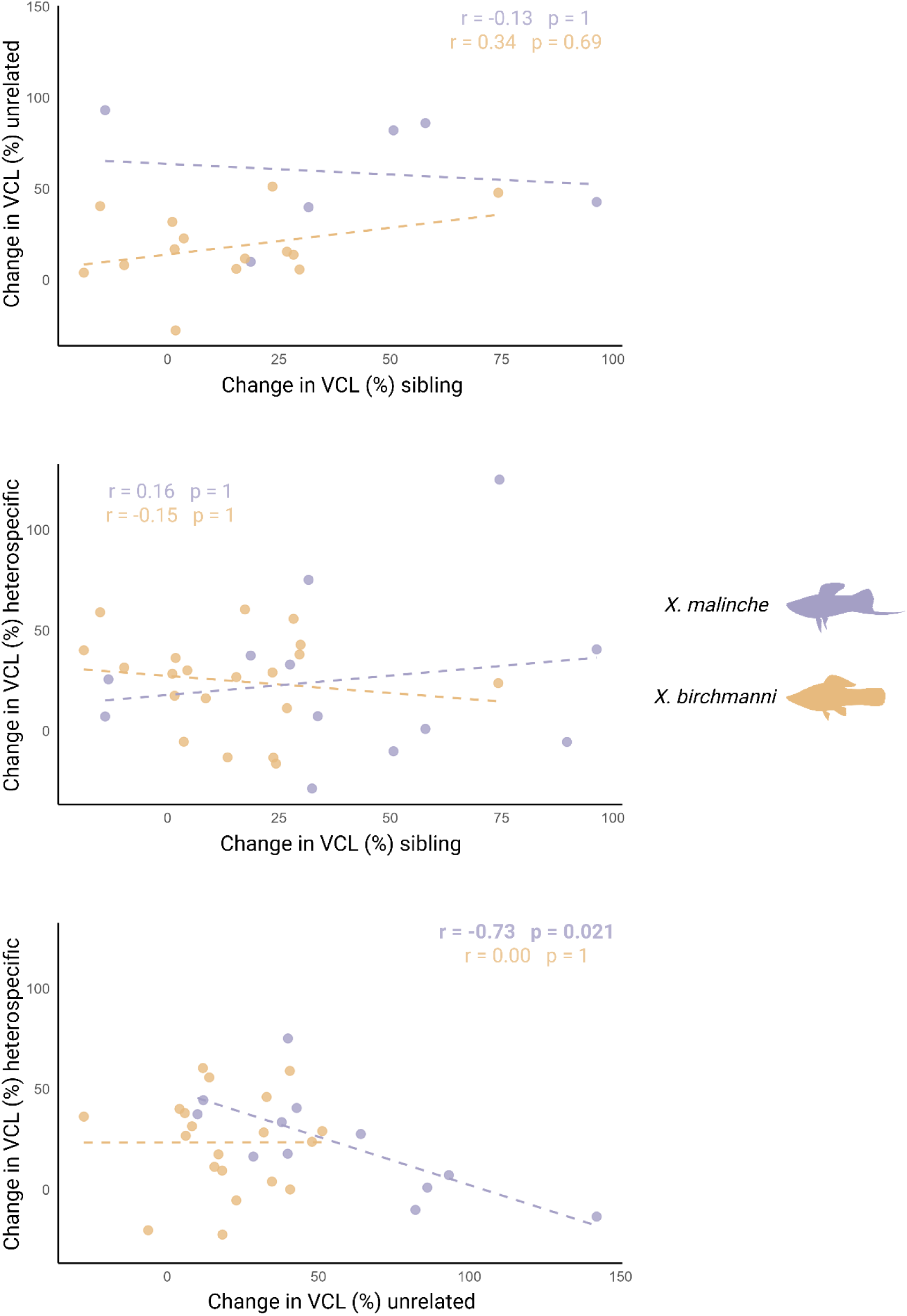
Covariation in sperm velocity responses across male types. Pearson correlations between FRF responses (change in sperm velocity in FRF relative to control) across the three sperm treatments (sibling, unrelated conspecific, and heterospecific) for *X. birchmanni* (orange) and *X. malinche* (purple). Each point represents a female’s effect on a male VCL change across the paired female environments. Correlations were computed separately for each pairwise comparison and species; p-values were adjusted for multiple comparisons using Holm’s correction.

There was no evidence for a trade-off between response to heterospecifics and siblings in either species (all adjusted p-values ≥ 0.69). In *X. birchmanni,* there was no correlation between responses to conspecific versus heterospecific sperm (r = 0, p_adj = 1) while, *X. malinche* showed a strong negative association between responses to unrelated conspecific and heterospecific males (r = −0.73, p_adj = 0.021). Females therefore vary in postmating choosiness (Rosenthal 2017), with some females both enhancing sperm velocity of conspecific males and reducing the velocity of heterospecific sperm.

## Discussion

Our results reveal pronounced and asymmetric effects of female reproductive fluid (FRF) on sperm performance between two naturally hybridizing sister species, indicating rapid divergence in postmating mechanisms of mate choice. In particular, FRF from *X. malinche* females selectively enhanced sperm velocity of conspecific males relative to heterospecifics, providing clear evidence of postmating discrimination against heterospecific sperm. By contrast, FRF from *X. birchmanni* females did not favor conspecific sperm over heterospecifics, revealing a striking species asymmetry in postmating isolation. This asymmetry was further supported by patterns of covariation in sperm responses: in *X. malinche*, female fluids that most strongly enhanced conspecific sperm performance also most strongly suppressed heterospecific sperm, whereas no such coupling was observed in *X. birchmanni*. Together, these results indicate fundamental differences between species in how FRF mediates postmating choice. Given the recent divergence of these sister species (46), our findings suggest that FRF-mediated postmating barriers can evolve rapidly.

The two species also diverged in how FRF interacted with intrinsic ejaculate performance. In *X. malinche*, conspecific FRF preferentially enhanced the velocity of lower-performing ejaculates, whereas in *X. birchmanni* sperm performance in conspecific and sibling fluids was reduced unless ejaculates were already fast. These patterns suggest that FRF-mediated selection acts in a phenotype-dependent manner, with species-specific consequences for fertilization dynamics. Conspecific preference in *X. malinche* is driven by a disproportionate velocity boost to slower sperm, presumably giving a fertilization advantage to conspecific sperm that would otherwise be outcompeted by heterospecifics. In guppies, FRF effects produce a similar boost for compatible but poor-ejaculate males (7, 10), suggesting that such compensatory FRF-mediated effects may be a broader feature of postmating female influence.

Evidence for inbreeding avoidance was weaker than for heterospecific discrimination, with no evidence of a trade-off between these two forms of postmating choice. While caution is warranted given limited power, the absence of a negative association argues against a simple mechanistic trade-off between avoiding sperm that are too genetically dissimilar and those that are too similar (9). Instead, these results raise the possibility that postmating female-mediated selection can simultaneously act along multiple axes of genetic compatibility, as in plants (19).

Mate choice emerges from the combined action of mechanisms operating before, during and after mating, and theory predicts that constraints on premating choice should favor the evolution of stronger postmating discrimination (8, 3, 47). In this context, the stronger FRF-mediated postmating response in *X. malinche* may compensate for reduced choosiness or limited opportunities for mate assessment prior to copulation. Across these sister species, the species characterized by weaker premating preferences exhibits more pronounced postmating effects on sperm performance, biasing fertilization outcomes toward unrelated conspecific males. This interpretation is consistent with known differences in sensory ecology and mating behavior between *X. malinche* and *X. birchmanni*. Whereas *X. birchmanni* females show robust, experience-dependent premating preferences for conspecifics, premating discrimination in *X. malinche* is weaker (30, 32, 27) and accompanied by behaviors that may limit mate sampling (33, 34). Under such conditions, postmating mechanisms provide an additional opportunity for discrimination after copulation, potentially buffering against mismatches that escape premating choice. Ecological differences between the species may further shape opportunities for postmating discrimination and help explain why FRF-mediated effects are stronger in *X. malinche*. This species inhabits small, spatially structured highland streams in which dispersal is limited, increasing the likelihood that siblings encounter one another as adults. In contrast, *X. birchmanni* occupies more connected lowland habitats where dispersal distances are greater (48). These differences alter the probability of encountering related mates, independent of the magnitude of inbreeding costs. The upstream–downstream structure of the river system also creates asymmetries in heterospecific encounters. Downstream populations of *X. birchmanni* may receive migrants from upstream *X. malinche*, whereas upstream populations are less likely to experience reciprocal movement. Such directional contact can generate unequal opportunities for hybridization between species. In this context, the stronger postmating discrimination observed in *X. malinche* is consistent with the idea that variation in encounter structure—both among relatives and between species—contributes to divergence in postmating mechanisms between sister taxa.

Female reproductive fluid favors conspecific males, but still permits the transit of heterospecific sperm. Indeed, these two species hybridize readily when premating barriers are disrupted. The conspecific bias of FRF is therefore only one of multiple prezygotic barriers to gene flow. While these barriers may have evolved through reinforcement after secondary contact (8, 3, 49), they are stronger in the upstream species where migration from downstream is rare (29). FRF has been proposed to be a rapidly evolvable trait, in part because some reproductive tract proteins may experience reduced pleiotropic constraint (17, 18). A plausible alternative to reinforcement is that FRF phenotypes have diverged in isolation due to their role in intraspecific mate choice, perhaps accelerated by sexual conflict with sperm and ejaculate traits (50, 51). The interspecific genetics of FRF in these two species, and the power to statistically detect incompatibilities (20, 21) offer a promising avenue for identifying these genetic interactions. Notwithstanding consistent preferences for conspecifics, there was substantial variation in FRF responses to individual males. In *X. malinche* in particular, an increased boost to conspecific sperm was correlated with a reduced one to heterospecifics: females thus vary in the strength of their FRF as a barrier to hybridization.

Future work should address proximate modulators of FRF activity: in particular, whether and how social context, such as the composition of available males, alters the selective effects exerted by FRF on sperm. In the *birchmanni-malinche* system, premating preferences are shaped by early (31, 33) and recent (52) social experience, as well as by recent nutritional condition (28), environmental stress (53), and age (54). Interestingly, there was no effect of fineness (a proxy for female nutritional condition and reproductive state, 41) nor standard length (a proxy for age and social dominance, 43) on FRF activity, suggesting that this variation is not simply explained by female condition or age, but may instead reflect discrimination among sperm phenotypes of individual males or compatibility effects at the level of individual gametes.

Our results align with mechanistic expectations for how female reproductive fluid (FRF) can shape sperm function. The influence of FRF likely reflects a combination of physical parameters (osmolality, pH, ion concentrations) that modulate motility and activation, together with biochemical constituents (sugars, amino acids, hormones and proteins) that affect sperm physiology (55-58). These interacting factors offer plausible pathways by which FRF could differentially enhance conspecific sperm and suppress heterospecific or closely related sperm, thereby creating opportunities for female-mediated selection on genetic compatibility. One possible mechanism underlying such compatibility effects is recognition mediated by hypervariable molecular cues. Under this scenario, disassortative postmating selection could arise through matched variation in peptide/protein products of genes such as the major histocompatibility complex (MHC), as well as odorant and taste receptors, conserved within species, if expressed in both sperm or ejaculate and female reproductive fluid (59, 27, 60, 10). A similar molecular mechanism could also explain the absence of an apparent trade-off between conspecific preference and inbreeding avoidance after mating, in contrast to before (9). In flowering plants, the same molecular system prevents selfing (extreme inbreeding) and interspecific pollination (19) through a shared SLR1–SRK receptor system that discriminates pollen identity and activates the same rejection pathway in cases of selfing or interspecific mismatch. An analogous system may mediate sperm-FRF interactions without imposing a functional tradeoff.

Studies of female reproductive fluid, and of postmating mechanisms more generally, have typically focused on the behavior of sperm in a female “environment”, with motile sperm and an invisible fluid playing the old roles of the active male and the passive female (61-63). But the female reproductive fluid, like the sensory and endocrine physiology that modulates premating choice, is an evolving and dynamic phenotype in its own right (10), that gives females agency over reproductive outcomes even when matings are hasty or coerced (64, 36). Differences between hybridizing sister species set the stage for genomic and functional studies (39, 65) of postmating mechanisms of mate choice and their evolutionary consequences.

## Author contribution

G.G.R.; C.G.; L.P.; and E.M. conceptualized the study and developed the methodology. L.P.; E.M. performed the investigation and curated the data. E.M.; L.P. and A.D. conducted formal analysis. O.C.D. and J.H.M. prepared the experimental families used in the study. L.P. and E.M. prepared the original draft of the manuscript, contributing equally, and all authors contributed to writing – review and editing. L.P. handled visualization, while G.G.R. supervised the project, managed project administration, and secured funding.

## Acknowledgements

We gratefully acknowledge the staff of the Centro de Investigaciones Científicas de las Huastecas ‘Aguazarca’ (CICHAZ) for their invaluable support and dedicated care of the fish used in this study.

## Supplementary Information

### Repeatability

The repeatability of sperm performance traits in the absence and presence of FRF was tested using the rptR package with a Gaussian distribution. Variables were log-transformed to meet the assumptions of a Gaussian distribution, and the analysis was based on 1,000 permutations. Sperm curvilinear velocity (VCL) was highly repeatable within individual males (Table 1 SI, Model 1 SI), both in absence and presence of FRF.

**Table 1 SI.**
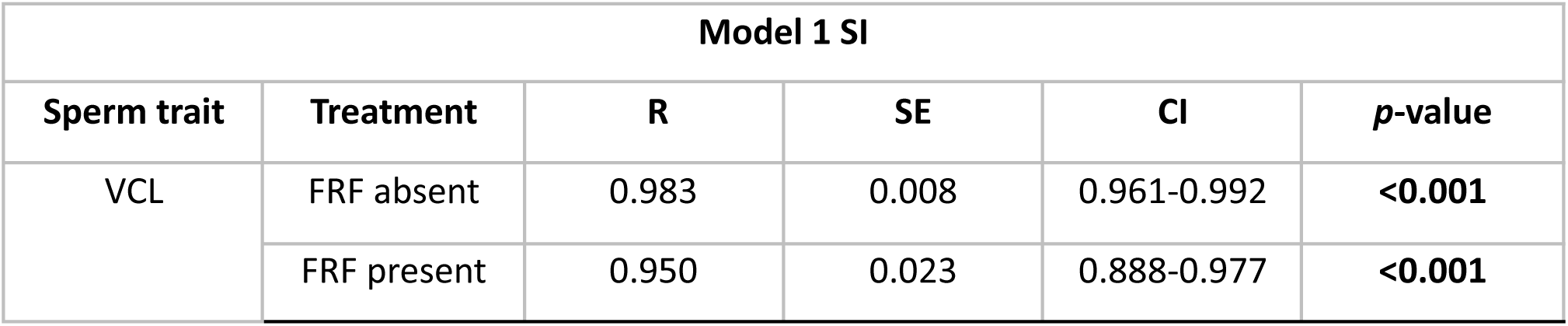
Repeatability test on sperm traits. VCL= Curvilinear Velocity; R=Repeatability; SE=Standard Error; CI=Confidence Interval.

### Interspecific differences in sperm performance

When two closely related species hybridize, ejaculate features are expected to overlap. To compare sperm performance in the control solution of the two closely related species *X.birchmanni* and *X.malinche*, we investigated mean VCL running a Welch two samples *t*-test. VCL mean is unequal across species (*t*-test Table 2; *p*=<0.001), specifically, *X. birchmanni* VCL is significantly higher than *X. malinche*.

**Table 2 SI.**
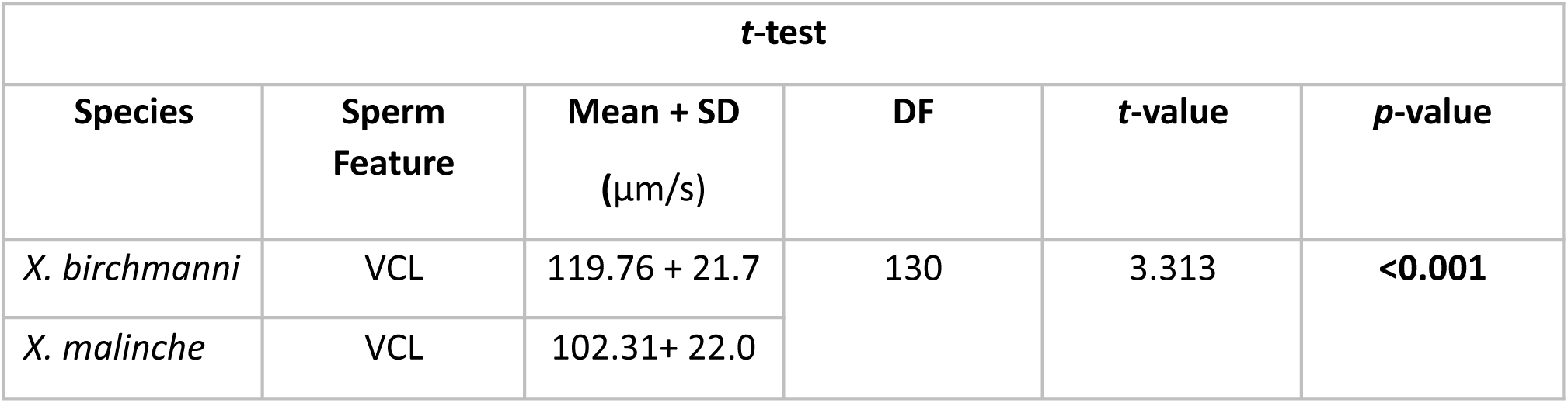
Welch two samples *t*-test on mean VCL in the control solution in *X. birchmanni* and *X. malinche*. N = 70 (39 *X.birchmanni*; 31 *X.malinche)*; Mean = mean VCL µm/s; DF = degrees of freedom. Significant terms are in bold.

Analysis of individual female trajectories revealed substantial variation in how reproductive fluid modulated sperm velocity across male types within each species. In *X. birchmanni*, female effects were relatively modest and consistent across sibling, unrelated conspecific and heterospecific males. In contrast, *X. malinche* females exhibited stronger and more heterogeneous modulation of sperm performance, with some individuals markedly enhancing velocity in unrelated or sibling conspecific sperm while showing reduced effects in heterospecific sperm. These patterns mirror the species-level results (Fig. S1) and indicate greater female-mediated modulation of sperm velocity in *X. malinche* compared to *X. birchmanni*.

**Figure S1.**
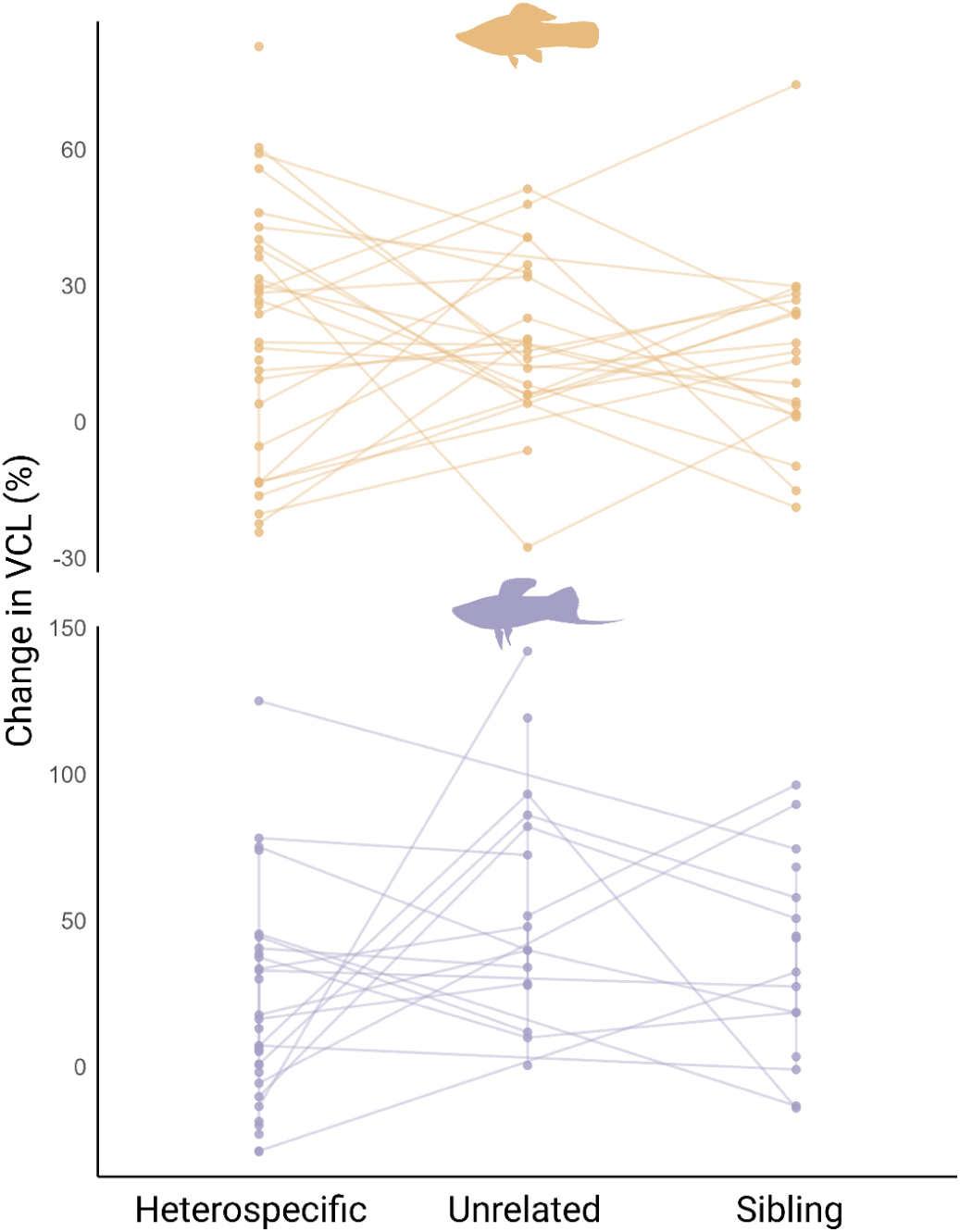
Individual variation in FRF-mediated changes in sperm velocity across male types. Individual trajectories of FRF effects on sperm velocity (ΔVCL) across the three male types - sibling, unrelated conspecific, and heterospecific- for *X. birchmanni* (top panel, orange) and *X. malinche* (bottom panel, purple). Each line represents a single female and connects the ΔVCL values measured for sperm from the three male types exposed to her reproductive fluid, illustrating within-species variation in female-mediated sperm responses.

To verify the robustness of Model 1 (Main Text), we sequentially removed non-significant predictors. First, male standard length (SL) was excluded (Model 2 SI; Table 3 SI), followed by the female fineness index (Model 3 SI; Table 4 SI). Model comparison tests (likelihood-ratio tests, p=0.123) showed no significant change in model fit after removing these terms, and the parameter estimates and significance levels of the remaining predictors were virtually unchanged. This confirms that the inclusion of male and female SL and fineness index did not influence the main results.

**Table 3 SI.**
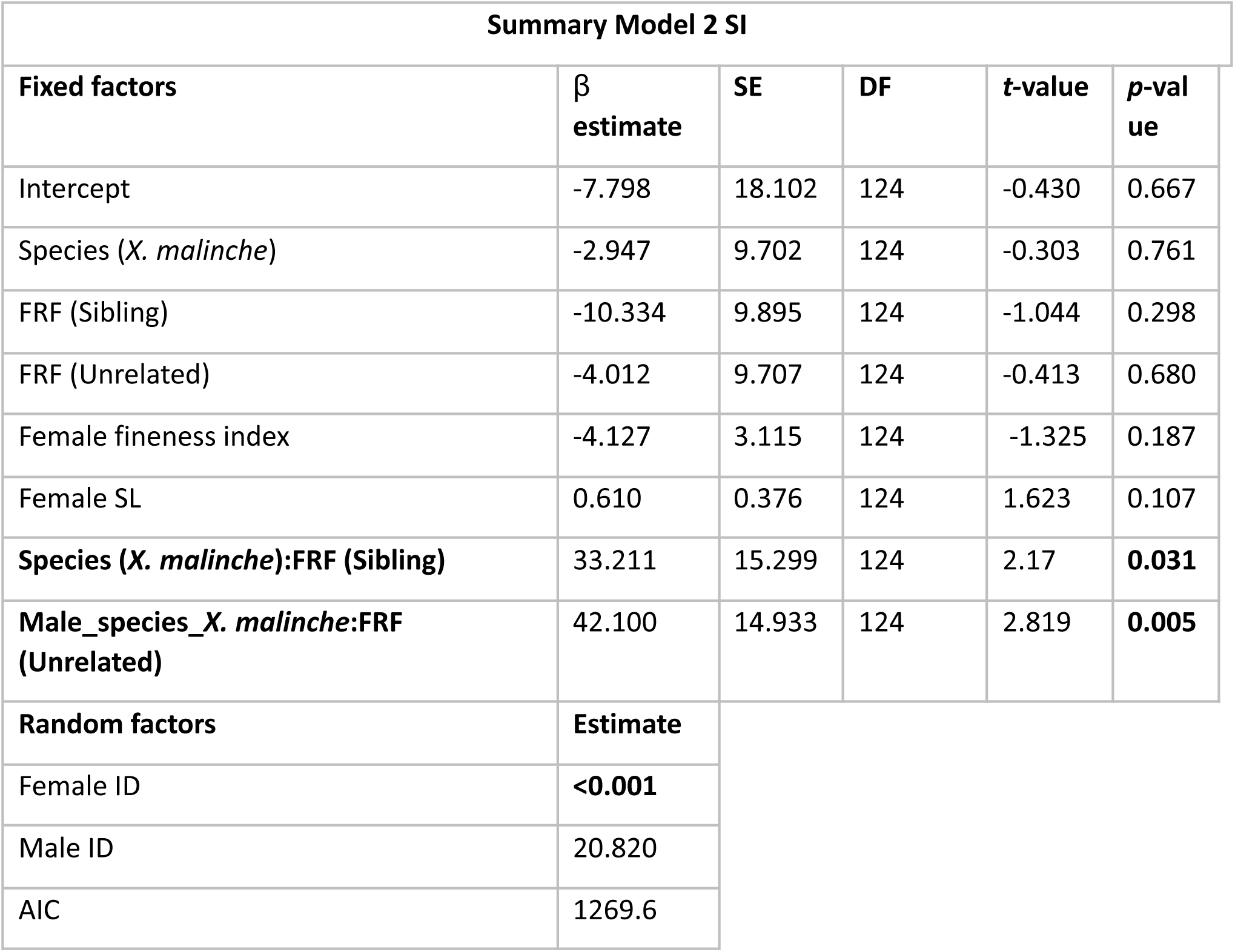
Results of LME including VCL change as dependent variable; fineness index, female SL, male species, FRF type, their interaction as fixed factors; and female and male identity as random factors. Significant terms are in bold. Regression coefficients of the interactions are referred to the *X. malinche* species while *X. birchmanni* was set to 0. FRF type was treated as a categorical variable with heterospecific as the reference level. SE=Standard Error; DF=Degrees of Freedom.

**Table 4 SI.**
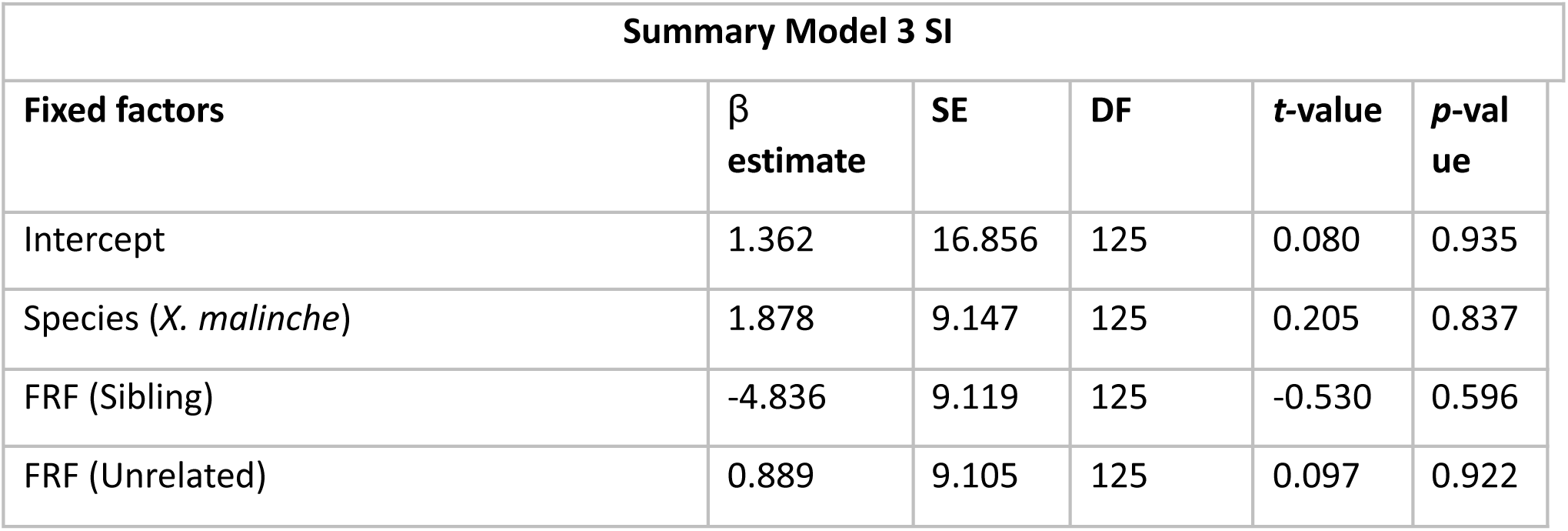
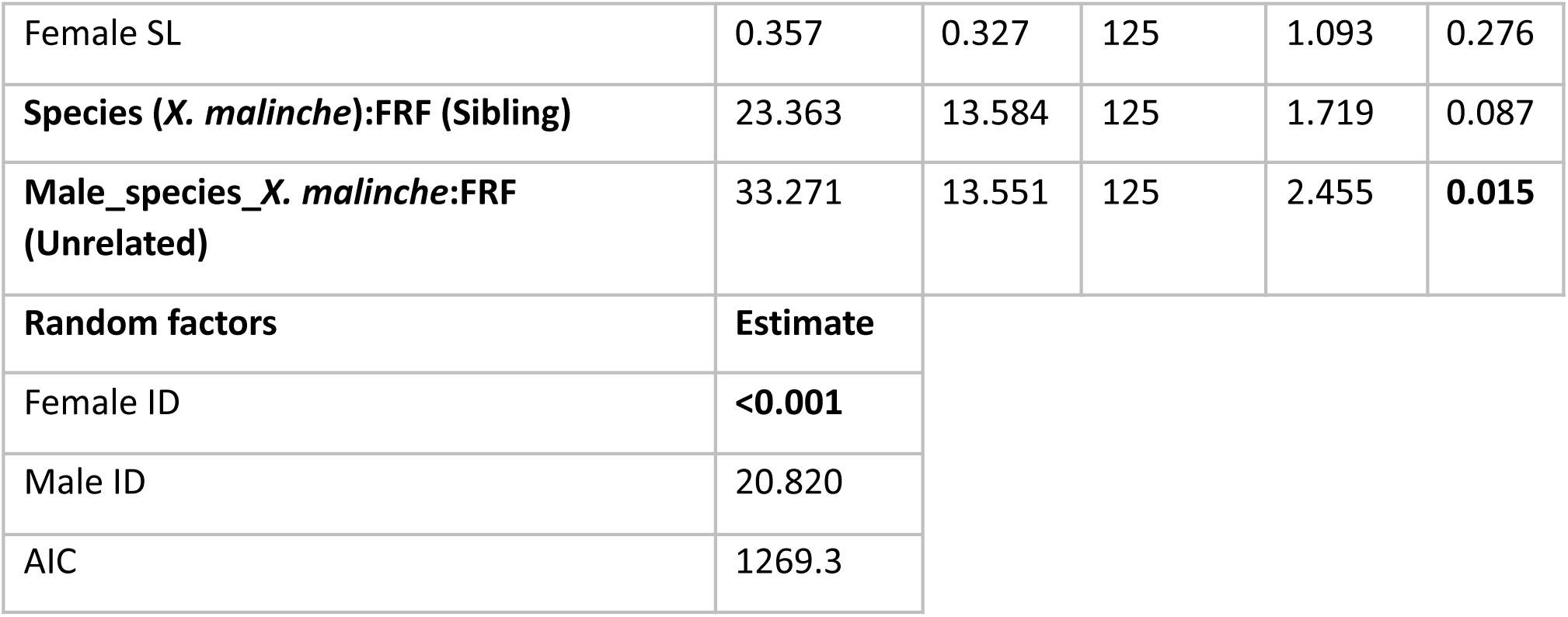
Results of LME including VCL change as dependent variable; female SL, male species, FRF type, their interaction as fixed factors; and female and male identity as random factors. Significant terms are in bold. Regression coefficients of the interactions are referred to the *X. malinche* species while *X. birchmanni* was set to 0. FRF type was treated as a categorical variable with heterospecific as the reference level. FRF=Female Reproductive Fluid; SE=Standard Error; DF=Degrees of Freedom.

**Table 5 SI.**
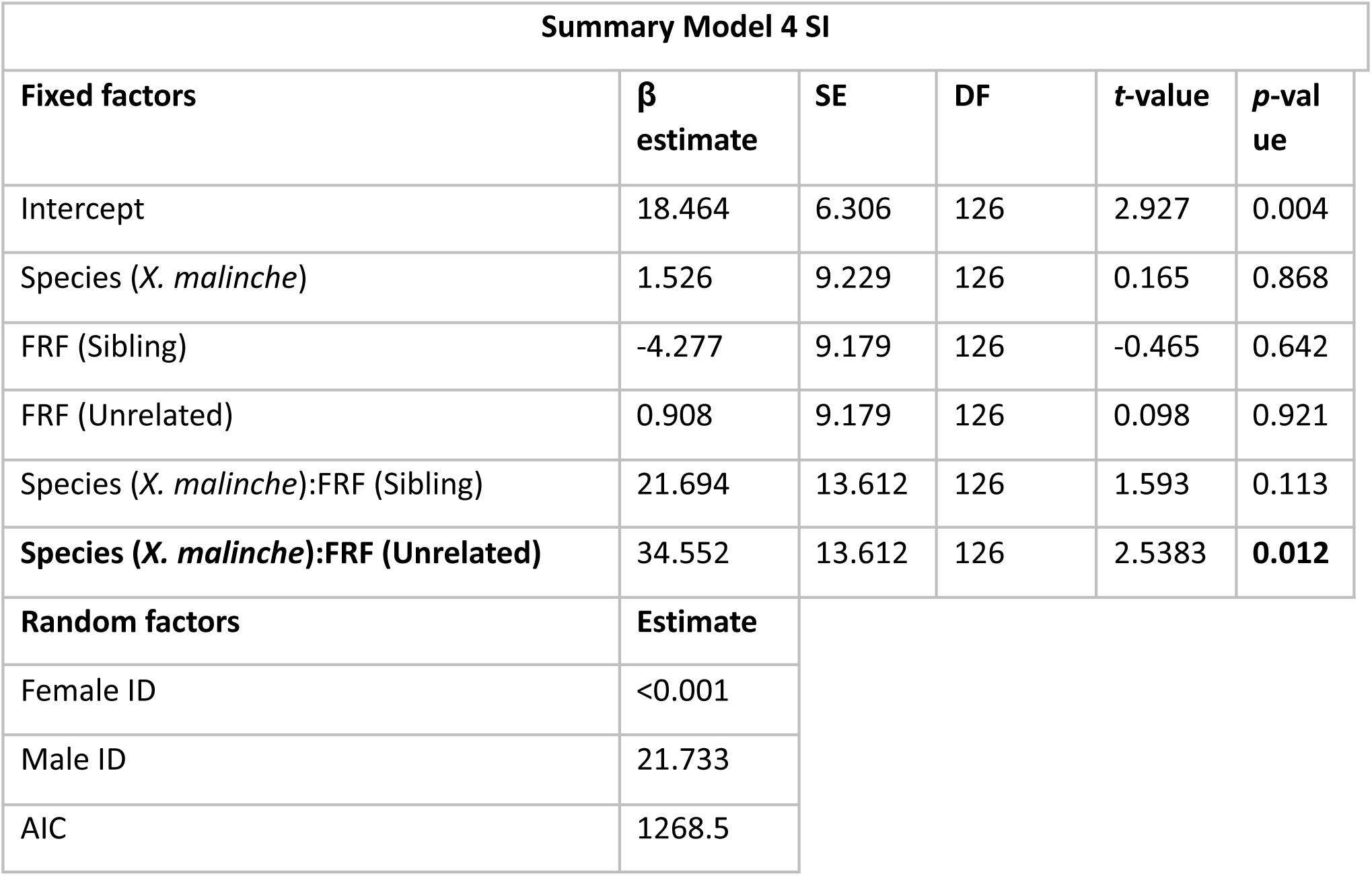
Results of LME including VCL change as dependent variable; male species, FRF type, their interaction as fixed factors, and female and male identity as random factors. Significant terms are in bold. Regression coefficients of the interactions are referred to the *X. malinche* species while *X. birchmanni* was set to 0. FRF type was treated as a categorical variable with heterospecific as the reference level. FRF=Female Reproductive Fluid; SE=Standard Error; DF=Degrees of Freedom.

We tested whether sperm performance differed among male and female genetic families, using a linear mixed-effects model with the percentage change in sperm velocity (VCL change) as the dependent variable. Male and female families were included as a fixed factor, and male and female identity were treated as random effects to account for repeated measures within individuals.

A likelihood-ratio test comparing models with and without the male family term revealed a highly significant improvement in model fit (χ² = 601.23, df = 14, *p* < 0.001), indicating strong family-level effects on sperm performance. Examination of the fixed effects showed that males from family AD20 exhibited a markedly higher VCL change than the reference family (β = 68.21 ± 24.01, *t* = 2.84, *p* = 0.006), while family AD4 showed a marginal increase (β = 27.64 ± 13.86, *t* = 1.99, *p* = 0.051). Differences among the remaining families were not statistically significant (all *p* > 0.07) (Table 6 SI).

Random-effect variances indicated substantial variation among males (Estimate = 15.6), but negligible variation among females (Estimate = <0.001), suggesting that most of the unexplained variance in sperm performance occurred at the male level. Overall, these results demonstrate that male genetic background significantly affects sperm velocity responses, supporting the view that sperm performance traits are at least partly heritable in poeciliids (66, 67).

**Table 6 SI.**
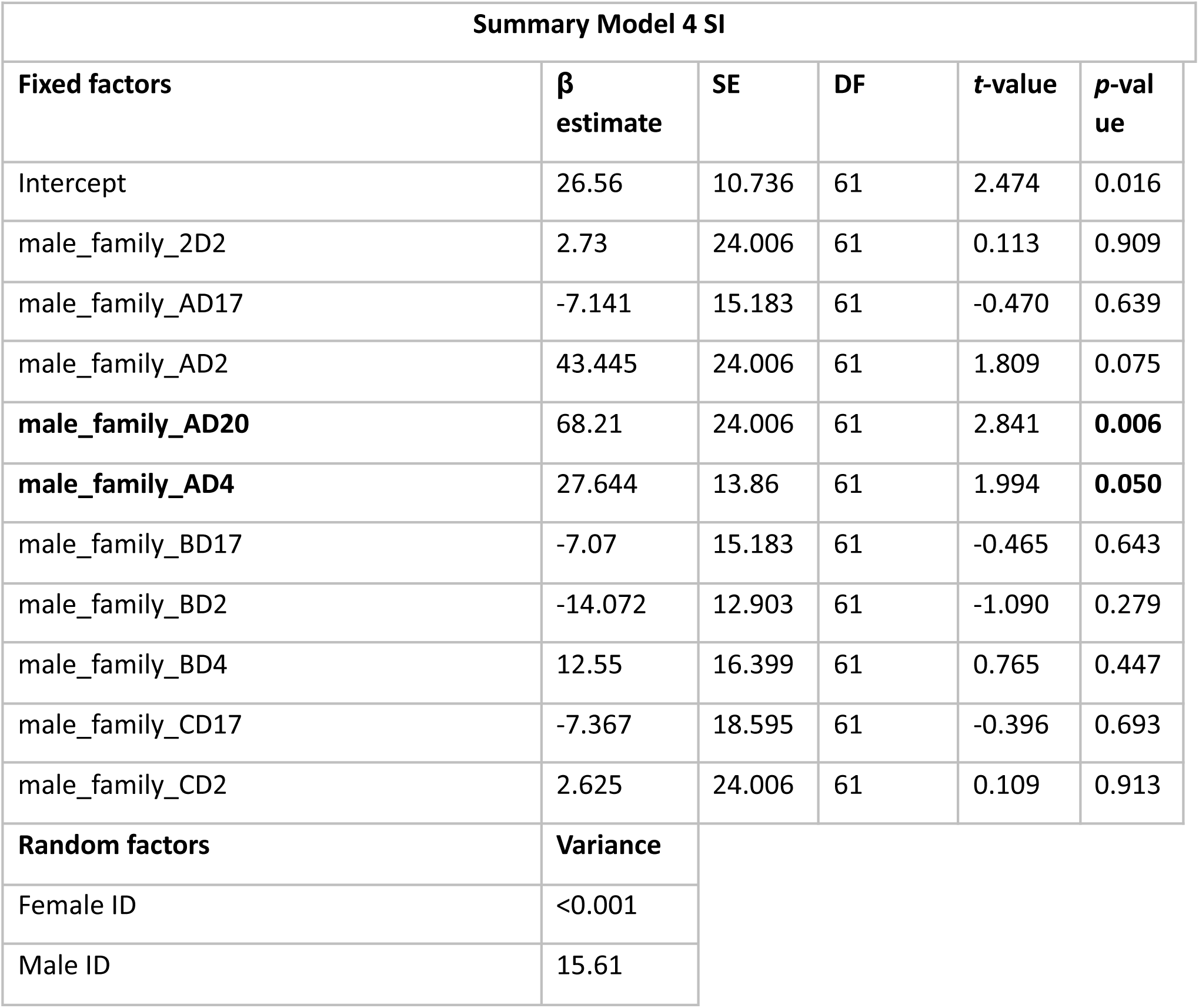
Results of LME including VCL change as dependent variable; male family, as fixed factor, and female and male identity as random factors. FRF type was treated as a categorical variable with heterospecific as the reference level. Significant terms are in bold. FRF=Female Reproductive Fluid; SE=Standard Error; DF=Degrees of Freedom.

To test whether sperm performance varied among female genetic families, we fitted a linear mixed-effects model with percentage change in sperm velocity (VCL change) as the dependent variable. Including female family as a fixed effect significantly improved the model compared with the reduced model lacking this term (likelihood-ratio test: χ² = 29.62, df = 12, p = 0.003), indicating significant family-level effects associated with the female genetic background.

Fixed effects showed that males mating with females from *birchmanni* family AD2 exhibited a significantly greater increase in VCL change compared to the reference family (β = 45.70 ± 18.81, t = 2.43, p = 0.018), and a similar positive effect was detected for *malinche* family AD20 (β = 43.71 ± 15.74, t = 2.78, p = 0.007). No statistically significant differences were detected among the remaining families (all p > 0.07). Random-effect estimates indicated substantial variance attributable to males (SD ≈ 14.7) (Table 7 SI).

Overall, these results demonstrate that female genetic background can also influence sperm velocity responses, although its contribution appears more limited than that observed for male family effects.

**Table 7 SI.**
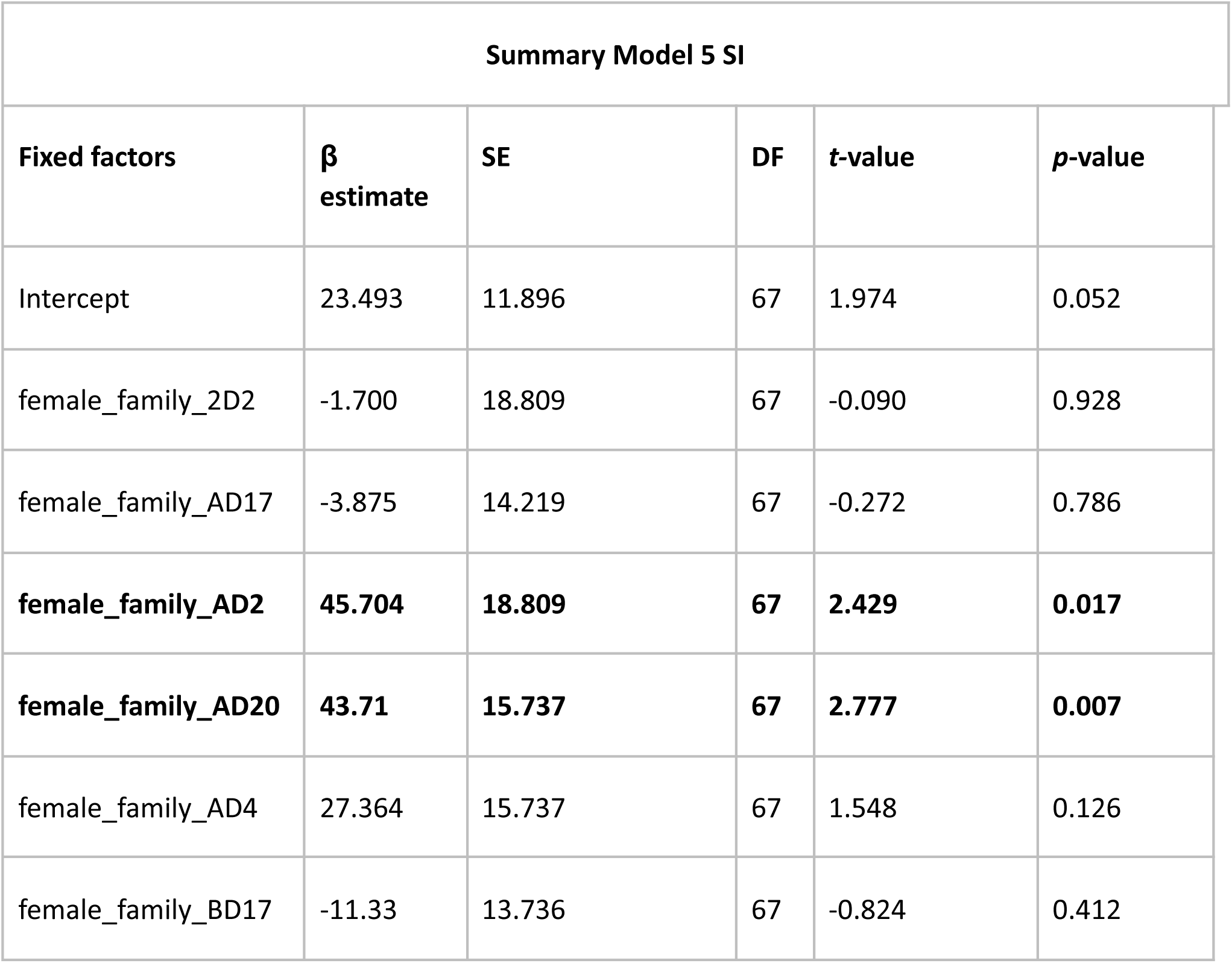
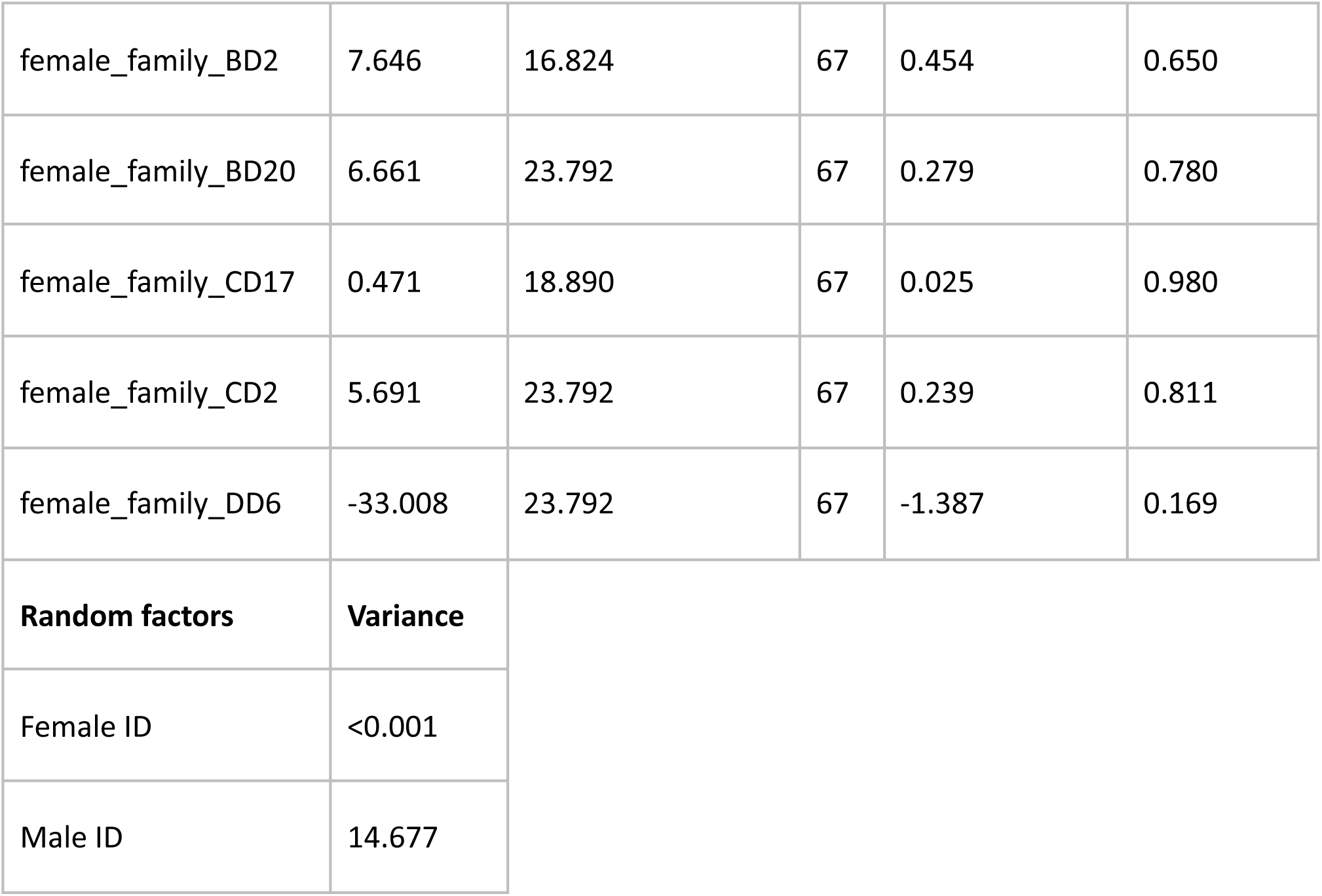
Results of LME including VCL change as dependent variable; female family, as fixed factor, and female and male identity as random factors. Significant terms are in bold. FRF=Female Reproductive Fluid; SE=Standard Error; DF=Degrees of Freedom.

### Fish history and husbandry

All procedures were approved under Texas A&M AUP 2020-0324. Fish used in these experiments were from outbred X. *birchmanni* (Coacuilco) and X. *malinche* (Chicayotla; see 53 for locality details) stock housed at CICHAZ descended from wild collections in 2019-2020 under Mexican federal Permiso de Pesca de Fomento no. PPF/DGOPA-002/19.

All work took place at the Centro de Investigaciones Cientìficas de las Huastecas “Aguazarca” (CICHAZ), Calnali, Hidalgo, Mexico, using municipal Río Calnali water dechlorinated with sodium thiosulfate and/or aerosolization in air. Subjects in this experiment were offspring of parents reared in outdoor mesocosm populations (concrete flow-through tanks and plastic recirculating tubs) operating under a natural photoperiod and ambient temperatures. Mesocosm fish foraged on naturally occurring invertebrates and periphyton and received a daily supplement of Ken’s Tropical Green Granules (up to 4.5 g per tank/day, approximately scaled by tank biomass). The fish tested in this experiment were reared indoors in rooms optimized for *X. birchmanni* (20-25 °C) and X. *malinche* (15-20 °C) husbandry. Indoor fish were fed twice daily on a rotating diet including the above as well as live Artemia salina nauplii and Repashy Soilent Green gel. Photoperiods were adjusted seasonally and were set to 11 h light∶13 h dark during these experiments.

### Generating experimental cohorts

To generate sibling cohorts for testing postmating inbreeding avoidance, we needed to isolate offspring of matings from one male and one female. Since poecilid females can store sperm for months (68), we generated virgin females by isolating 15 juveniles (<1 cm standard length) collected from large stock populations and placed into 2,000 L single-species outdoor mesocosms (7 per species). Mature male poeciliids develop a modified intromittent organ, the gonopodium, derived from the anterior portion of the anal fin. We inspected fish visually every 2-3 days and removed any individuals that showed signs of a thickening anal fin or other male sexually-dimorphic traits. The remaining individuals were allowed to grow and mate with a single conspecific male from a different stock population to minimize the risk of inbreeding depression in the sibling cohorts. We thus generated 9 sets of full siblings nested within 3 half-sibling *X. birchmanni* groups and 6 sets of full siblings nested within 3 half-sibling *X. malinche* groups (Table 8 and 9 SI). 2-12 full siblings were housed in mixed sex groups in 60 liter aquaria or in 90 liter aquaria subdivided into 4 compartments.

**Table 8 SI.**
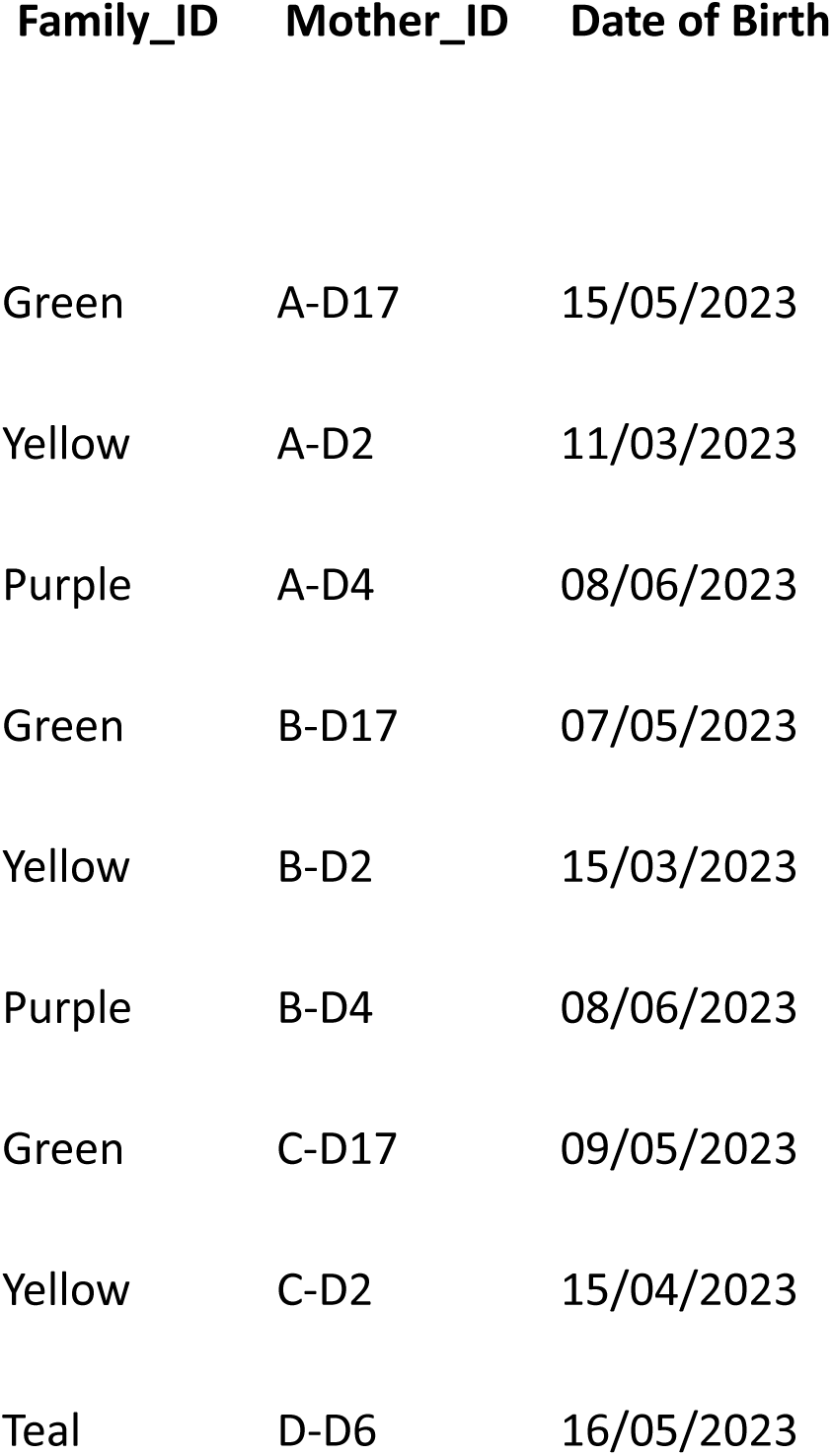
Families of ***Xiphophorus birchmanni*** included in this study. For each family, we report the family identifier (Family_ID), the maternal identifier (Mother_ID), and the date of birth of the offspring. Dates are reported in day/month/year format. Cells marked as NA indicate missing or unrecorded birth dates.

**Table 9.**
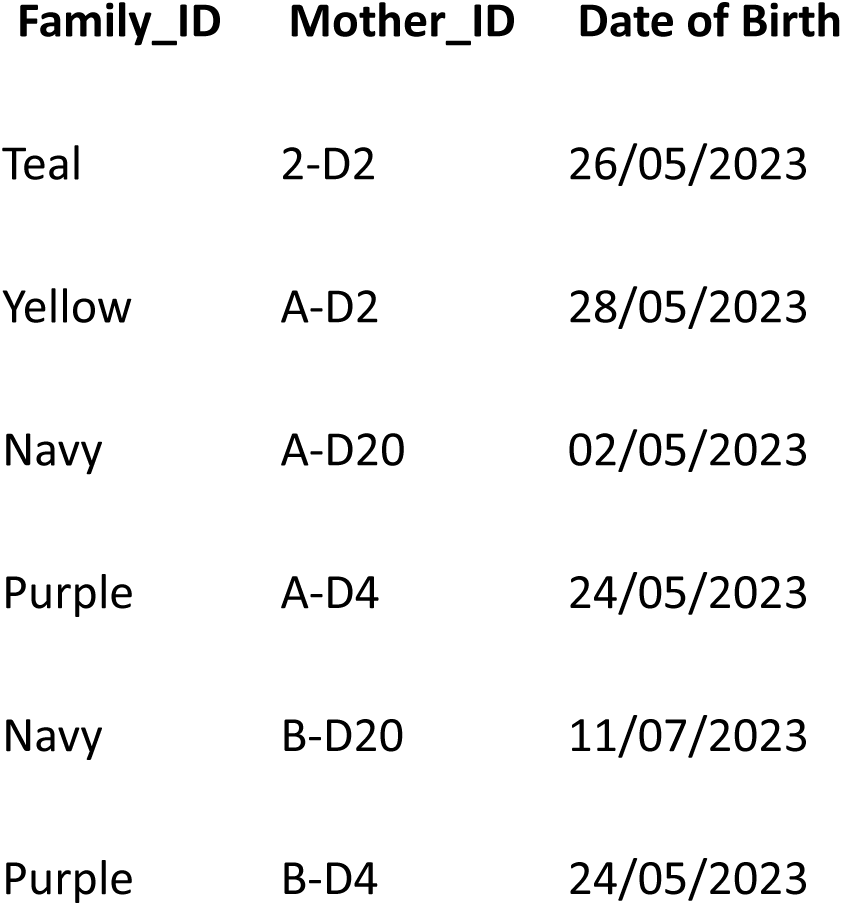
Families of ***Xiphophorus malinche*** included in this study. For each family, we report the family identifier (Family_ID), the maternal identifier (Mother_ID), and the date of birth of the offspring. Dates are reported in day/month/year format.

All fish were sexually mature and dimorphic, indicating reproductive condition at the time of the experiment; *X. birchmanni* individuals ranged from 222 to 327 days of age, whereas *X. malinche* individuals ranged from 157 to 275 days of age. Most males produced an ejaculate upon stripping, although 18 males did not release sperm and were therefore excluded from sperm assays. Fish were housed by full-sibling cohort in recirculating semi-closed systems using Calnali river water from the municipal system, dechlorinated through aeration and application of sodium thiosulfate. During rearing, *X. malinche* and *X. birchmanni* were housed in separate, adjacent, temperature-optimized fishrooms that tracked seasonal variation in highland (*X.malinche*) and lowland (*X.birchmanni*) temperatures. Prior to starting the experiment, fish were gradually acclimatized to a common, intermediate temperature of 22°C, well within the normal thermal range of both species (69).

## Notes

https://doi.org/10.5281/zenodo.20163079

### Competing Interest Statement

The authors have declared no competing interest.

## References

1. Edmands S, 2007. Between a rock and a hard place: evaluating the relative risks of inbreeding and outbreeding for conservation and management. Molecular ecology, 16(3), 463–475.

2. Eberhard WG, 1966. Female control: sexual selection by cryptic female choice. Princeton University Press.

3. Firman RC, Gasparini C, Manier MK, Pizzari T, 2017. Postmating female control: 20 years of cryptic female choice. Trends in Ecology & Evolution 32, 368–382.

4. Garlovsky MD, Whittington E, Albrecht T, Arenas-Castro H, Castillo DM, Keais GL, Larson EL, Moyle LC, Plakke M, Reifová R, Snook RR, Ålund M, Weber AAT, 2024. Synthesis and scope of the role of postmating prezygotic isolation in speciation. Cold Spring Harbor Perspectives in Biology 16, a041429.

5. Charlesworth D, Vekemans X, Castric V, Glemin S, 2005. Plant self-incompatibility systems: a molecular evolutionary perspective. New Phytologist 168, 61–69.

6. Cutter AD, 2019. Reproductive transitions in plants and animals: selfing syndrome, sexual selection and speciation. New Phytologist 224, 1080–1094.

7. Gasparini C, Pilastro A, 2011. Cryptic female preference for genetically unrelated males is mediated by ovarian fluid in the guppy. Proceedings of the Royal Society B: Biological Sciences 278, 2495–2501.

8. Birkhead TR, Pizzari T, 2002. Postcopulatory sexual selection. Nature Reviews Genetics 3, 262–273.

9. Dorsey OC, Rosenthal GG, 2023. A taste for the familiar: explaining the inbreeding paradox. Trends in Ecology & Evolution 38, 132–142.

10. Gasparini C, Pilastro A, Evans JP, 2020. The role of female reproductive fluid in sperm competition. Philosophical Transactions of the Royal Society B 375, 20200077.

11. Miller RL, Mojares JJ, Ram JL, 1994. Species-specific sperm attraction in the zebra mussel (Dreissena polymorpha) and the quagga mussel (Dreissena bugensis). Canadian Journal of Zoology 72, 1764–1770.

12. Riffell JA, Krug PJ, Zimmer RK, 2004. The ecological and evolutionary consequences of sperm chemoattraction. Proceedings of the National Academy of Sciences of the United States of America 101, 4501–4506.

13. Yeates SE, Diamond SE, Einum S, Emerson BC, Holt WV, Gage MJG, 2013. Cryptic choice of conspecific sperm controlled by the impact of ovarian fluid on sperm swimming behavior. Evolution 67, 3523–3536.

14. Myers J, Bradford A, Hallas V, Lawson L, Pitcher T, Dunham R, Butts I, 2020. Channel catfish ovarian fluid differentially enhances blue catfish sperm performance. Theriogenology 149, 62–71.

15. Cramer ERA, Ålund M, McFarlane SE, Johnsen A, Qvarnström A, 2016a. Females discriminate against heterospecific sperm in a natural hybrid zone. Evolution 70, 1844–1855.

16. Cramer ER, Stensrud E, Marthinsen G, Hogner S, Johannessen LE, Laskemoen T, Eybert MC, Slagsvold T, Lifjeld JT, Johnsen A, 2016b. Sperm performance in conspecific and heterospecific female fluid. Ecology and Evolution 6, 1363–1377.

17. Beirão J, Purchase CF, Wringe BF, Fleming IA, 2015. Inter-population ovarian fluid variation differentially modulates sperm motility in Atlantic cod *Gadus morhua*. Journal of Fish Biology 87, 54–68.

18. Devigili A, Fitzpatrick JL, Gasparini C, Ramnarine IW, Pilastro A, Evans JP, 2018. Possible glimpses into early speciation: The effect of ovarian fluid on sperm velocity accords with post-copulatory isolation between two guppy populations. Journal of Evolutionary Biology 31, 66–74.

19. Liu B, Li M, Qiu J et al., 2024. A pollen selection system links self and interspecific incompatibility in the Brassicaceae. Nature Ecology & Evolution 8, 1129–1139.

20. Schumer M, Cui R, Powell DL, Dresner R, Rosenthal GG, Andolfatto P, 2014. High-resolution mapping reveals hundreds of genetic incompatibilities in hybridizing fish species. eLife 3, e02535.

21. Schumer M, Brandvain Y, 2016. Determining epistatic selection in admixed populations. Molecular Ecology 25, 2577–2591.

22. Powell DL, García-Olazábal M, Keegan M, Reilly P, Du K, Díaz-Loyo AP, Banerjee S, Blakkan D, Reich D, Andolfatto P, Rosenthal GG, Schartl M, Schumer M, 2020. Natural hybridization reveals incompatible alleles that cause melanoma in swordtail fish. Science, 368(6492), 731–736.

23. Moran BM, Payne CY, Powell DL, Iverson EN, Donny AE, Banerjee SM, et al., 2024. A lethal mitonuclear incompatibility in complex I of natural hybrids. Nature 626, 119–127.

24. Paczolt KA, Passow CN, Delclos PJ, Kindsvater HK, Jones AG, Rosenthal GG, 2015. Multiple mating and reproductive skew in parental and introgressed females of the live-bearing fish Xiphophorus birchmanni. Journal of Heredity 106, 57–66.

25. Squire MK, 2015. Mate choice and multiple paternity in the Xiphophorus malinche/X. birchmanni hybrid system. Doctoral dissertation.

26. McLennan DA, Ryan MJ, 1999. Interspecific recognition and discrimination based upon olfactory cues in northern swordtails. Evolution 53, 880–888.

27. Cui R, Delclos PJ, Schumer M, Rosenthal GG, 2017. Early social learning triggers neurogenomic expression changes in a swordtail fish. Proceedings of the Royal Society B: Biological Sciences 284.

28. Fisher HS, Wong B, Rosenthal GG, 2006. Alteration of the chemical environment disrupts communication in a freshwater fish. Proceedings of the Royal Society B: Biological Sciences 273, 1187–1193.

29. Moran BM, Ramírez-Duarte WF, Powell DL, Yang TTL, Gunn TR, Jofre-Rodríguez GI, Payne CY, Iverson ENK, Preising GA, Banerjee SM, et al. 2025. Increased rates of hybridization in swordtail fish are associated with water pollution. bioRxiv. doi: 10.1101/2025.04.22.649978.

30. Wong BB, Rosenthal GG, 2006. Female disdain for swords in a swordtail fish. The American Naturalist 167, 136–140.

31. Verzijden MN, Rosenthal GG, 2011. Effects of sensory modality on learned mate preferences in female swordtails. Animal Behaviour 82, 557–562.

32. Verzijden MN, Ten Cate C, Servedio MR, Kozak GM, Boughman JW, Svensson EI, 2012. The impact of learning on sexual selection and speciation. Trends in Ecology & Evolution 27, 511–519.

33. Delclos PJ, Forero SA, Rosenthal GG, 2020. Divergent neurogenomic responses shape social learning of personality and mate preference. Journal of Experimental Biology 223, jeb220707.

34. Johnson JB, Culumber ZW, Easterling R, Rosenthal GG, 2015. Boldness and predator evasion in naturally hybridizing swordtails. Current Zoology 61, 596–603.

35. Gage MJG, Macfarlane CP, Yeates S, Ward RG, Searle JB, Parker GA, 2004. Spermatozoal traits and sperm competition in Atlantic salmon. Current Biology 14, 44–47.

36. Pinzoni L, Rasotto MB, Gasparini C, 2024. Sperm performance in the race for fertilization: the influence of female reproductive fluid. Royal Society Open Science 11.

37. Boschetto C, Gasparini C, Pilastro A, 2011. Sperm number and velocity affect sperm competition success in the guppy (Poecilia reticulata). Behavioral Ecology and Sociobiology 65, 813–821.

38. Gasparini C, Simmons LW, Beveridge M, Evans JP, 2010. Sperm swimming velocity predicts competitive fertilization success in the green swordtail Xiphophorus helleri. PloS one, 5(8), e12146.

39. Powell DL, Payne C, Banerjee SM, Keegan M, Bashkirova E, Cui R, et al., 2021. The genetic architecture of variation in the sexually selected sword ornament. Current Biology 31, 923–935.

40. Harper C, Lawrence C, 2016. The laboratory zebrafish. CRC Press.

41. Fisher HS, Rosenthal GG, 2006. Hungry females show stronger mating preferences. Behavioral Ecology 17, 979–981.

42. Potter H, Kramer CR, 2000. Ultrastructural observations on sperm storage in Xiphophorus maculatus. Journal of Morphology 245, 110–129.

43. Ramsey ME, Wong RY, Cummings ME, 2011. Estradiol, reproductive cycle and preference behavior in a northern swordtail. General and Comparative Endocrinology 170, 381–390.

44. Cohen J, 1988. Statistical power for the behavioral sciences. Lawrence Erlbaum.

45. Evans JP, 2011. Patterns of genetic variation and covariation in ejaculate traits. Heredity 106, 869–875.

46. Schumer M, Rosenthal GG, Andolfatto P, 2018. What do we mean when we talk about hybrid speciation? Heredity 120, 379–382.

47. Lüpold S, Tomkins JL, Simmons LW, Fitzpatrick JL, 2014. Female monopolization mediates the relationship between pre- and postcopulatory traits. Nature Communications 5, 3184.

48. Culumber ZW, Fisher HS, Tobler M, Mateos M, Barber PH, Sorenson MD, Rosenthal GG, 2011. Replicated hybrid zones of Xiphophorus swordtails along an elevational gradient. Molecular Ecology 20, 342–356.

49. Servedio MR, Noor MA, 2003. The role of reinforcement in speciation. Annual Review of Ecology, Evolution, and Systematics 34, 339–364.

50. Holland B, Rice WR, 1999. Experimental removal of sexual selection reverses intersexual antagonistic coevolution. Proceedings of the National Academy of Sciences 96, 5083–5088.

51. Gavrilets S, 2000. Rapid evolution of reproductive barriers driven by sexual conflict. Nature 403, 886–889.

52. Willis PM, Ryan MJ, Rosenthal GG, 2011. Encounter rates with conspecific males influence female mate choice. Behavioral Ecology 22, 1234–1240.

53. Willis PM, Rosenthal GG, Ryan MJ, 2012. Predation risk counteracts female preference for conspecifics. PLoS ONE 7, e34802.

54. Tudor MS, Morris MR, 2009. Experience and female preference for symmetry. Ethology 115, 812–822.

55. Teves ME, Barbano F, Guidobaldi HA, Sanchez R, Miska W, Giojalas LC, 2006. Progesterone at the picomolar range is a chemoattractant for mammalian spermatozoa. Fertility and Sterility 86, 745–749.

56. Kekäläinen J, Larma I, Linden M, Evans JP, 2015. Lectin staining and flow cytometry reveals female-induced sperm acrosome reaction and surface carbohydrate reorganization. Scientific Reports 5, 15321.

57. Kholodnyy V, Gadêlha H, Cosson J, Boryshpolets S, 2019. How do freshwater fish sperm find the egg? The physicochemical factors guiding the gamete encounters of externally fertilizing freshwater fish. Reviews in Aquaculture 12, 1165–1192.

58. Pitnick S, Wolfner MF, Dorus S, 2020. Postejaculatory modifications to sperm (PEMS). Biological Reviews 95, 365–392.

59. Ziegler A, Santos PSC, Kellermann T, Uchanska-Ziegler B, 2010. Self/nonself perception and the extended MHC. Self/Nonself 1, 176–191.

60. Kekäläinen J, Evans JP, 2017. Female-induced regulation of sperm physiology. Evolution 71, 238–248.

61. Tang-Martínez Z, 2016. Rethinking Bateman’s principles. The Journal of Sex Research 53, 532–559.

62. Perry JC, Rowe L, 2018. Sexual conflict in its ecological setting. Philosophical Transactions of the Royal Society B 373.

63. Rosenthal GG, Ryan MJ, 2022. Sexual selection and the ascent of women. Science 375, eabi6308.

64. Brennan PL, Prum RO, 2015. Genital coevolution and sexual conflict. Cold Spring Harbor Perspectives in Biology 7, a017749.

65. Payne C, Bovio R, Powell DL, Gunn TR, Banerjee SM, Grant V, et al., 2024. Genomic insights into thermotolerance in hybridizing swordtails. Molecular Ecology 33, e16489.

## References

66. Gasparini C, Devigili A, Dosselli R, Pilastro A, 2013. Pattern of inbreeding depression, condition dependence, and additive genetic variance in Trinidadian guppy ejaculate traits. Ecology and evolution, 3(15), 4940–4953.

67. Evans JP, 2010. Quantitative genetic evidence that males trade attractiveness for ejaculate quality in guppies. Proceedings of the Royal Society B: Biological Sciences, 277(1697), 3195–3201.

68. López-Sepulcre A, Gordon SP, Paterson IG, Bentzen P, Reznick DN, 2013. Beyond lifetime reproductive success: the posthumous reproductive dynamics of male Trinidadian guppies. Proc. R. Soc. B 280 (1763).

69. Culumber ZW, Shepard DB, Coleman SW, Rosenthal GG, Tobler M, 2012. Physiological adaptation along environmental gradients and replicated hybrid zone structure in swordtails (Teleostei: *Xiphophorus*). J. Evol. Biol. 25, 1800–1814.

