## Supplementary Information for "Female reproductive fluid evolves rapidly to favor conspecific sperm"

*Repeatability*

The repeatability of sperm performance traits in the absence and presence of FRF was tested using the rptR package with a Gaussian distribution. Variables were log-transformed to meet the assumptions of a Gaussian distribution, and the analysis was based on 1,000 permutations. Sperm curvilinear velocity (VCL) was highly repeatable within individual males (Table 1 SI, Model 1 SI), both in absence and presence of FRF.

Table 1 SI. Repeatability test on sperm traits. VCL= Curvilinear Velocity; R=Repeatability; SE=Standard Error; CI=Confidence Interval.

| **Model 1 SI** | | | | | |
|---|---|---|---|---|---|
| **Sperm trait** | **Treatment** | **R** | **SE** | **CI** | ***p*-value** |
| VCL | FRF absent | 0.983 | 0.008 | 0.961-0.992 | **<0.001** |
| | FRF present | 0.950 | 0.023 | 0.888-0.977 | **<0.001** |

*Interspecific differences in sperm performance*

When two closely related species hybridize, ejaculate features are expected to overlap. To compare sperm performance in the control solution of the two closely related species *X.birchmanni* and *X.malinche*, we investigated mean VCL running a Welch two samples *t*-test. VCL mean is unequal across species (*t*-test Table 2; *p*=<0.001), specifically, *X. birchmanni* VCL is significantly higher than *X. malinche*.

Table 2 SI. Welch two samples *t*-test on mean VCL in the control solution in *X. birchmanni* and *X. malinche*. N = 70 (39 *X.birchmanni*; 31 *X.malinche)*; Mean = mean VCL µm/s; DF = degrees of freedom. Significant terms are in bold.

| ***t*-test** |
|---|

| Species | Sperm Feature | Mean + SD (µm/s) | DF | *t*-value | *p*-value |
|---|---|---|---|---|---|
| *X. birchmanni* | VCL | 119.76 + 21.7 | 130 | 3.313 | **<0.001** |
| *X. malinche* | VCL | 102.31+ 22.0 | | | |

Analysis of individual female trajectories revealed substantial variation in how reproductive fluid modulated sperm velocity across male types within each species. In *X. birchmanni*, female effects were relatively modest and consistent across sibling, unrelated conspecific and heterospecific males. In contrast, *X. malinche* females exhibited stronger and more heterogeneous modulation of sperm performance, with some individuals markedly enhancing velocity in unrelated or sibling conspecific sperm while showing reduced effects in heterospecific sperm. These patterns mirror the species-level results (Fig. S1) and indicate greater female-mediated modulation of sperm velocity in *X. malinche* compared to *X. birchmanni*.

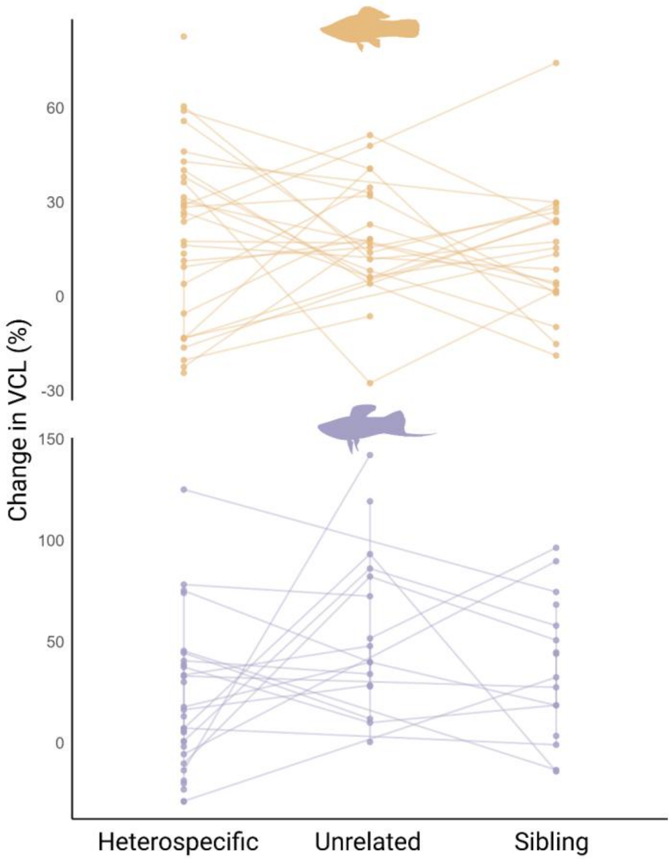


**Figure S1. Individual variation in FRF-mediated changes in sperm velocity across male types.** Individual trajectories of FRF effects on sperm velocity (ΔVCL) across the three male types - sibling , unrelated conspecific, and heterospecific- for *X. birchmanni* (top panel, orange) and *X. malinche* (bottom panel, purple). Each line represents a single female and connects the ΔVCL values measured for sperm from the three male types exposed to her reproductive fluid, illustrating within-species variation in female-mediated sperm responses.

To verify the robustness of Model 1 (Main Text), we sequentially removed non-significant predictors. First, male standard length (SL) was excluded (Model 2 SI; Table 3 SI), followed by the female fineness index (Model 3 SI; Table 4 SI). Model comparison tests (likelihood-ratio tests, p=0.123) showed no significant change in model fit after removing these terms, and the parameter estimates and significance levels of the remaining predictors were virtually unchanged. This confirms that the inclusion of male and female SL and fineness index did not influence the main results.

Table 3 SI. Results of LME including VCL change as dependent variable; fineness index, female SL, male species, FRF type, their interaction as fixed factors; and female and male identity as random factors. Significant terms are in bold. Regression coefficients of the interactions are referred to the *X. malinche* species while *X. birchmanni* was set to 0. FRF type was treated as a categorical variable with heterospecific as the reference level.  SE=Standard Error; DF=Degrees of Freedom.

| **Summary Model 2 SI** | | | | | |
|---|---|---|---|---|---|
| **Fixed factors** | **β estimate** | **SE** | **DF** | ***t*-value** | ***p*-value** |
| Intercept | -7.798 | 18.102 | 124 | -0.430 | 0.667 |
| Species (*X. malinche*) | -2.947 | 9.702 | 124 | -0.303 | 0.761 |
| FRF (Sibling) | -10.334 | 9.895 | 124 | -1.044 | 0.298 |
| FRF (Unrelated) | -4.012 | 9.707 | 124 | -0.413 | 0.680 |
| Female fineness index | -4.127 | 3.115 | 124 | -1.325 | 0.187 |
| Female SL | 0.610 | 0.376 | 124 | 1.623 | 0.107 |
| **Species (*X. malinche*):FRF (Sibling)** | 33.211 | 15.299 | 124 | 2.17 | **0.031** |
| **Male_species_*X. malinche*:FRF (Unrelated)** | 42.100 | 14.933 | 124 | 2.819 | **0.005** |
| **Random factors** | **Estimate** | | | | |
| Female ID | **<0.001** | | | | |
| Male ID | 20.820 | | | | |
| AIC | 1269.6 | | | | |

Table 4 SI. Results of LME including VCL change as dependent variable; female SL, male species, FRF type, their interaction as fixed factors; and female and male identity as random factors. Significant terms are in bold. Regression coefficients of the interactions are referred to the *X. malinche* species while *X. birchmanni* was set to 0. FRF type was treated as a categorical variable with heterospecific as the reference level. FRF=Female Reproductive Fluid; SE=Standard Error; DF=Degrees of Freedom.

| **Summary Model 3 SI** |
|---|

| **Fixed factors** | **β estimate** | **SE** | **DF** | ***t*-value** | ***p*-value** |
|---|---|---|---|---|---|
| Intercept | 1.362 | 16.856 | 125 | 0.080 | 0.935 |
| Species (*X. malinche*) | 1.878 | 9.147 | 125 | 0.205 | 0.837 |
| FRF (Sibling) | -4.836 | 9.119 | 125 | -0.530 | 0.596 |
| FRF (Unrelated) | 0.889 | 9.105 | 125 | 0.097 | 0.922 |
| Female SL | 0.357 | 0.327 | 125 | 1.093 | 0.276 |
| **Species (*X. malinche*):FRF (Sibling)** | 23.363 | 13.584 | 125 | 1.719 | 0.087 |
| **Male_species_*X. malinche*:FRF (Unrelated)** | 33.271 | 13.551 | 125 | 2.455 | **0.015** |
| **Random factors** | **Estimate** | | | | |
| Female ID | **<0.001** | | | | |
| Male ID | 20.820 | | | | |
| AIC | 1269.3 | | | | |

Table 5 SI. Results of LME including VCL change as dependent variable; male species, FRF type, their interaction as fixed factors, and female and male identity as random factors. Significant terms are in bold. Regression coefficients of the interactions are referred to the *X. malinche* species while *X. birchmanni* was set to 0. FRF type was treated as a categorical variable with heterospecific as the reference level. FRF=Female Reproductive Fluid; SE=Standard Error; DF=Degrees of Freedom.

| **Summary Model 4 SI** | | | | | |
|---|---|---|---|---|---|
| **Fixed factors** | **β estimate** | **SE** | **DF** | ***t*-value** | ***p*-value** |
| Intercept | 18.464 | 6.306 | 126 | 2.927 | 0.004 |
| Species (*X. malinche*) | 1.526 | 9.229 | 126 | 0.165 | 0.868 |
| FRF (Sibling) | -4.277 | 9.179 | 126 | -0.465 | 0.642 |
| FRF (Unrelated) | 0.908 | 9.179 | 126 | 0.098 | 0.921 |
| Species (*X. malinche*):FRF (Sibling) | 21.694 | 13.612 | 126 | 1.593 | 0.113 |
| **Species (*X. malinche*):FRF (Unrelated)** | 34.552 | 13.612 | 126 | 2.5383 | **0.012** |
| **Random factors** | **Estimate** | | | | |
| Female ID | <0.001 | | | | |

| | |
|---|---|
| Male ID | 21.733 |
| AIC | 1268.5 |

We tested whether sperm performance differed among male and female genetic families, using a linear mixed-effects model with the percentage change in sperm velocity (VCL change) as the dependent variable. Male and female families were included as a fixed factor, and male and female identity were treated as random effects to account for repeated measures within individuals.
A likelihood-ratio test comparing models with and without the male family term revealed a highly significant improvement in model fit ($\chi^2 = 601.23$, df = 14, $p < 0.001$), indicating strong family-level effects on sperm performance. Examination of the fixed effects showed that males from family AD20 exhibited a markedly higher VCL change than the reference family ($\beta = 68.21 \pm 24.01$, $t = 2.84$, $p = 0.006$), while family AD4 showed a marginal increase ($\beta = 27.64 \pm 13.86$, $t = 1.99$, $p = 0.051$). Differences among the remaining families were not statistically significant (all $p > 0.07$) (Table 6 SI). Random-effect variances indicated substantial variation among males (Estimate = 15.6), but negligible variation among females (Estimate = <0.001), suggesting that most of the unexplained variance in sperm performance occurred at the male level.
Overall, these results demonstrate that male genetic background significantly affects sperm velocity responses, supporting the view that sperm performance traits are at least partly heritable in poeciliids (66, 67).

Table 6 SI. Results of LME including VCL change as dependent variable; male family, as fixed factor, and female and male identity as random factors. FRF type was treated as a categorical variable with heterospecific as the reference level. Significant terms are in bold. FRF=Female Reproductive Fluid; SE=Standard Error; DF=Degrees of Freedom.

| **Summary Model 4 SI** | | | | | |
|---|---|---|---|---|---|
| **Fixed factors** | **β estimate** | **SE** | **DF** | ***t*-value** | ***p*-value** |
| Intercept | 26.56 | 10.736 | 61 | 2.474 | 0.016 |
| male_family_2D2 | 2.73 | 24.006 | 61 | 0.113 | 0.909 |
| male_family_AD17 | -7.141 | 15.183 | 61 | -0.470 | 0.639 |
| male_family_AD2 | 43.445 | 24.006 | 61 | 1.809 | 0.075 |
| **male_family_AD20** | 68.21 | 24.006 | 61 | 2.841 | **0.006** |
| **male_family_AD4** | 27.644 | 13.86 | 61 | 1.994 | **0.050** |
| male_family_BD17 | -7.07 | 15.183 | 61 | -0.465 | 0.643 |
| male_family_BD2 | -14.072 | 12.903 | 61 | -1.090 | 0.279 |
| male_family_BD4 | 12.55 | 16.399 | 61 | 0.765 | 0.447 |

| | | | | | |
|---|---|---|---|---|---|
| male_family_CD17 | -7.367 | 18.595 | 61 | -0.396 | 0.693 |
| male_family_CD2 | 2.625 | 24.006 | 61 | 0.109 | 0.913 |
| **Random factors** | **Variance** | | | | |
| Female ID | <0.001 | | | | |
| Male ID | 15.61 | | | | |

To test whether sperm performance varied among female genetic families, we fitted a linear mixed-effects model with percentage change in sperm velocity (VCL change) as the dependent variable. Including female family as a fixed effect significantly improved the model compared with the reduced model lacking this term (likelihood-ratio test: $\chi^2 = 29.62$, df = 12, p = 0.003), indicating significant family-level effects associated with the female genetic background.

Fixed effects showed that males mating with females from *birchmanni* family AD2 exhibited a significantly greater increase in VCL change compared to the reference family (β = 45.70 ± 18.81, t = 2.43, p = 0.018), and a similar positive effect was detected for *malinche* family AD20 (β = 43.71 ± 15.74, t = 2.78, p = 0.007). No statistically significant differences were detected among the remaining families (all p > 0.07). Random-effect estimates indicated substantial variance attributable to males (SD ≈ 14.7) (Table 7 SI).

Overall, these results demonstrate that female genetic background can also influence sperm velocity responses, although its contribution appears more limited than that observed for male family effects.

Table 7 SI. Results of LME including VCL change as dependent variable; female family, as fixed factor, and female and male identity as random factors. Significant terms are in bold. FRF=Female Reproductive Fluid; SE=Standard Error; DF=Degrees of Freedom.

| **Summary Model 5 SI** | | | | | |
|---|---|---|---|---|---|
| **Fixed factors** | **β estimate** | **SE** | **DF** | ***t*-value** | ***p*-value** |
| Intercept | 23.493 | 11.896 | 67 | 1.974 | 0.052 |
| female_family_2D2 | -1.700 | 18.809 | 67 | -0.090 | 0.928 |
| female_family_AD17 | -3.875 | 14.219 | 67 | -0.272 | 0.786 |
| **female_family_AD2** | **45.704** | **18.809** | **67** | **2.429** | **0.017** |
| **female_family_AD20** | **43.71** | **15.737** | **67** | **2.777** | **0.007** |

| | | | | | |
|---|---|---|---|---|---|
| female_family_AD4 | 27.364 | 15.737 | 67 | 1.548 | 0.126 |
| female_family_BD17 | -11.33 | 13.736 | 67 | -0.824 | 0.412 |
| female_family_BD2 | 7.646 | 16.824 | 67 | 0.454 | 0.650 |
| female_family_BD20 | 6.661 | 23.792 | 67 | 0.279 | 0.780 |
| female_family_CD17 | 0.471 | 18.890 | 67 | 0.025 | 0.980 |
| female_family_CD2 | 5.691 | 23.792 | 67 | 0.239 | 0.811 |
| female_family_DD6 | -33.008 | 23.792 | 67 | -1.387 | 0.169 |
| **Random factors** | **Variance** | | | | |
| Female ID | <0.001 | | | | |
| Male ID | 14.677 | | | | |

Table 8 SI. Families of ***Xiphophorus birchmanni*** included in this study. For each family, we report the family identifier (Family_ID), the maternal identifier (Mother_ID), and the date of birth of the offspring. Dates are reported in day/month/year format. Cells marked as NA indicate missing or unrecorded birth dates.

| **Family_ID** | **Mother_ID** | **Date of Birth** |
|---|---|---|
| Green | A-D17 | 15/05/2023 |
| Yellow | A-D2 | 11/03/2023 |
| Purple | A-D4 | 08/06/2023 |
| Green | B-D17 | 07/05/2023 |
| Yellow | B-D2 | 15/03/2023 |
| Purple | B-D4 | 08/06/2023 |
| Green | C-D17 | 09/05/2023 |

| | | |
|---|---|---|
| Yellow | C-D2 | 15/04/2023 |
| Teal | D-D6 | 16/05/2023 |

Table 9. Families of ***Xiphophorus malinche*** included in this study. For each family, we report the family identifier (Family_ID), the maternal identifier (Mother_ID), and the date of birth of the offspring. Dates are reported in day/month/year format.

| **Family_ID** | **Mother_ID** | **Date of Birth** |
|---|---|---|
| Teal | 2-D2 | 26/05/2023 |
| Yellow | A-D2 | 28/05/2023 |
| Navy | A-D20 | 02/05/2023 |
| Purple | A-D4 | 24/05/2023 |
| Navy | B-D20 | 11/07/2023 |
| Purple | B-D4 | 24/05/2023 |

All fish were sexually mature and dimorphic, indicating reproductive condition at the time of the experiment; *X. birchmanni* individuals ranged from 222 to 327 days of age, whereas *X. malinche* individuals ranged from 157 to 275 days of age. Most males produced an ejaculate upon stripping, although 18 males did not release sperm and were therefore excluded from sperm assays. Fish were housed by full-sibling cohort in recirculating semi-closed systems using Calnali river water from the municipal system, dechlorinated through aeration and application of sodium thiosulfate. During rearing, *X. malinche* and *X. birchmanni* were housed in separate, adjacent, temperature-optimized fishrooms that tracked seasonal variation in highland (*X.malinche*) and lowland (*X.birchmanni*) temperatures. Prior to starting the experiment, fish were gradually acclimatized to a common, intermediate temperature of 22°C, well within the normal thermal range of both species (69).

### References

66. Gasparini C, Devigili A, Dosselli R, Pilastro A, 2013. Pattern of inbreeding depression, condition dependence, and additive genetic variance in Trinidadian guppy ejaculate traits. Ecology and evolution, 3(15), 4940-4953.

67. Evans JP, 2010. Quantitative genetic evidence that males trade attractiveness for ejaculate quality in guppies. Proceedings of the Royal Society B: Biological Sciences, 277(1697), 3195-3201.

68. López-Sepulcre A, Gordon SP, Paterson IG, Bentzen P, Reznick DN, 2013. Beyond lifetime reproductive success: the posthumous reproductive dynamics of male Trinidadian guppies. Proc. R. Soc. B 280 (1763).

69. Culumber ZW, Shepard DB, Coleman SW, Rosenthal GG, Tobler M, 2012. Physiological adaptation along environmental gradients and replicated hybrid zone structure in swordtails (Teleostei: *Xiphophorus*). J. Evol. Biol. 25, 1800–1814.
